# Ion channel-enhancing nanoparticles reprogram T cells in vivo for cancer immunotherapy

**DOI:** 10.64898/2026.11.16.767140

**Authors:** Ameet A. Chimote, Marat V. Khodoun, Anjali Iyer, Abdulaziz O. Alshwimi, Shweta Swain, Hannah S. Newton, Benjamin H. Hinrichs, Laura Conforti

## Abstract

In head and neck squamous cell carcinoma (HNSCC), immunotherapy is frequently ineffective due to the impaired ability of tumor-infiltrating T cells to sustain calcium-dependent cytotoxic effector functions. Kv1.3 potassium channels in T cells are essential for maintaining membrane potential, which is necessary for Ca^2+^ influx; however, they are suppressed within the tumor microenvironment, and no pharmacologic Kv1.3 activators are currently available. To address this limitation, we developed lipid nanoparticles (LNPs) functionalized with anti-CD5 antibodies to specifically deliver mRNA encoding Kv1.3 (KCNA3) to T cells, thereby increasing channel expression and function. Treatment of human T cells with KCNA3-LNPs resulted in increased Kv1.3 channel protein levels and activity, leading to enhanced Ca^2+^ influx and elevated secretion of effector cytokines (IL-2, IL-17, TNF-α, and IFN-γ) and cytotoxic molecules (Granzyme A, Perforin, Granulysin, and Fas ligand). In a human peripheral blood mononuclear cell-engrafted HNSCC humanized mouse model, peritumoral delivery of KCNA3-LNPs significantly reduced tumor burden and increased the infiltration of CD8⁺ Granzyme B⁺ cytotoxic T cells within the tumor, indicating improved antitumor immunity. Our study demonstrates that in vivo reprogramming of T cells through targeted Kv1.3 enhancement can overcome immunosuppressive barriers within the tumor microenvironment and enhance immune-mediated tumor clearance in vivo. This novel approach offers a scalable and cost-effective alternative to ex vivo T cell engineering and holds promise as a therapeutic strategy to improve immunotherapy outcomes in HNSCC and other solid tumors characterized by Kv1.3-dependent immune dysfunctions.

**One Sentence Summary:** Targeted mRNA delivery restores Kv1.3 channels in T cells, thereby reviving antitumor immunity and reducing head and neck tumor growth.

## INTRODUCTION

Head and neck squamous cell carcinoma (HNSCC) is an aggressive cancer with high mortality, particularly in high-risk cases where the 5-year overall survival rate remains below 50%. The debilitating comorbidities associated with the current standard of care, which includes surgery, radiation, and/or chemotherapy, highlight the urgent need for more effective treatment strategies (*1*). Monoclonal antibodies targeting the immune checkpoint receptor programmed cell death 1 (PD1) have been approved as first-line therapy for patients with metastatic or unresectable recurrent HNSCC, and more recently, for resectable locally advanced tumors expressing programmed cell death 1 ligand (PDL1). These therapies have demonstrated promising improvements in outcomes, including durable remission after therapy discontinuation (*2, 3*). However, 80–90% of patients either do not respond or relapse (*4*), underscoring the need for novel strategies to enhance immune-mediated antitumor responses and improve immunotherapy efficacy.

Immunosurveillance and the efficacy of immunotherapies in cancer rely on T lymphocytes and their ability to infiltrate tumors and eliminate cancer cells. Ion channels are critical regulators of these effector functions. T cells express ion channels that control Ca^2+^ signaling, which is required for cytotoxicity, cytokine secretion, and chemotaxis. Specifically, Kv1.3 and KCa3.1 potassium channels maintain the membrane potential that drives Ca^2+^ entry through Ca^2+^ channels and blockade of these channels suppresses T cell function (*5–11*). We have demonstrated that dysregulation of these channels contributes to failed immune surveillance in cancer and resistance to immunotherapy. Kv1.3, the focus of this study, is inhibited by hypoxia in the tumor microenvironment (TME) (*12*) and is functionally suppressed in tumor infiltrating T lymphocytes from patients with HNSCC, contributing to reduced cytotoxicity and cytokine release (*13, 14*).

Recently, we found that neoadjuvant treatment of HNSCC patients with a PD1-blocking antibody (anti-PD1) prior to surgery increased both KCa3.1 and Kv1.3 channel activity, accompanied by characteristic changes in patients who exhibited tumor response (*13*). Enhanced Kv1.3 activity and Ca^2+^ flux were observed in circulating cytotoxic T cells exclusively in responder patients, whereas KCa3.1 activity increased in both responders and non-responders. Elevated Kv1.3 function and Ca^2+^ signaling suggest that circulating T cells in responder patients may be in a more activated state and thus better poised to rapidly attack and kill cancer cells. Notably, these functional gains persisted long after tumor resection, indicating a mechanism that may contribute to the durable remission observed following anti-PD1 therapy (*13*). These observations support the concept that therapeutically increasing Kv1.3 activity could enhance T cell function, improve responses to standard immunotherapies, and potentially convert anti-PD1-resistant patients into responders. Although no selective activators of Kv1.3 are currently available, augmenting Kv1.3 expression offers a viable strategy. Adoptive transfer of Kv1.3-overexpressing T cells increased interleukin-2 (IL-2) and interferon-γ (IFN-γ) production, reduced tumor burden, and prolonged survival in a melanoma mouse model (*15*).

Here, we developed a strategy to selectively enhance the cytotoxic function and TME resistance of T cells in vivo without requiring ex vivo genetic editing. We engineered a liposomal mRNA formulation to drive overexpression of the Kv1.3 channel specifically within T cells. Lipid nanoparticles (LNPs) were used to encapsulate and protect the mRNA, facilitating efficient cellular delivery by overcoming barriers related to the molecule’s negative charge and hydrophilicity, which impede passive membrane diffusion (*16*). To further improve in vivo stability and enable cell-specific targeting, LNPs were functionalized with surface modifications. This specificity minimizes off-target effects by sparing other cells that rely on Kv1.3 for their function, such as monocytes/macrophages, platelets, adipose cells, and cancer cells (*17–19*). This strategy offers several advantages over traditional ex vivo T cell engineering, including reduced cost, simplified logistics, and the ability to transiently enhance Kv1.3 expression. The transient nature of mRNA expression, owing to its cytoplasmic localization, inherent instability, and dilution through cell division, limits prolonged activation, thereby reducing the risk of immune-related adverse events commonly associated with conventional cancer immunotherapies (*20*).

## RESULTS

### Association of high Kv1.3 gene (KCNA3) expression with improved survival in cancer patients

Kaplan–Meier survival analysis of HNSCC patients from The Cancer Genome Atlas (TCGA) cohort demonstrated that elevated expression of the Kv1.3 gene (KCNA3) correlates with improved overall survival (Fig. 1). Patients in the lower 50% of KCNA3 expression exhibited significantly reduced survival compared to those in the upper 50% (Cox coefficient = –0.264; P = 0.000524). A comparable trend was observed in clear cell renal cell carcinoma (ccRCC, Fig. S1), further supporting the relevance of Kv1.3 in cancer. These findings provide a strong rationale for strategies aimed at enhancing Kv1.3 expression in T cells. To this end, we developed LNPs for the targeted delivery of Kv1.3 mRNA to T cells, hereafter referred to as KCNA3-LNPs.

**Fig. 1.**
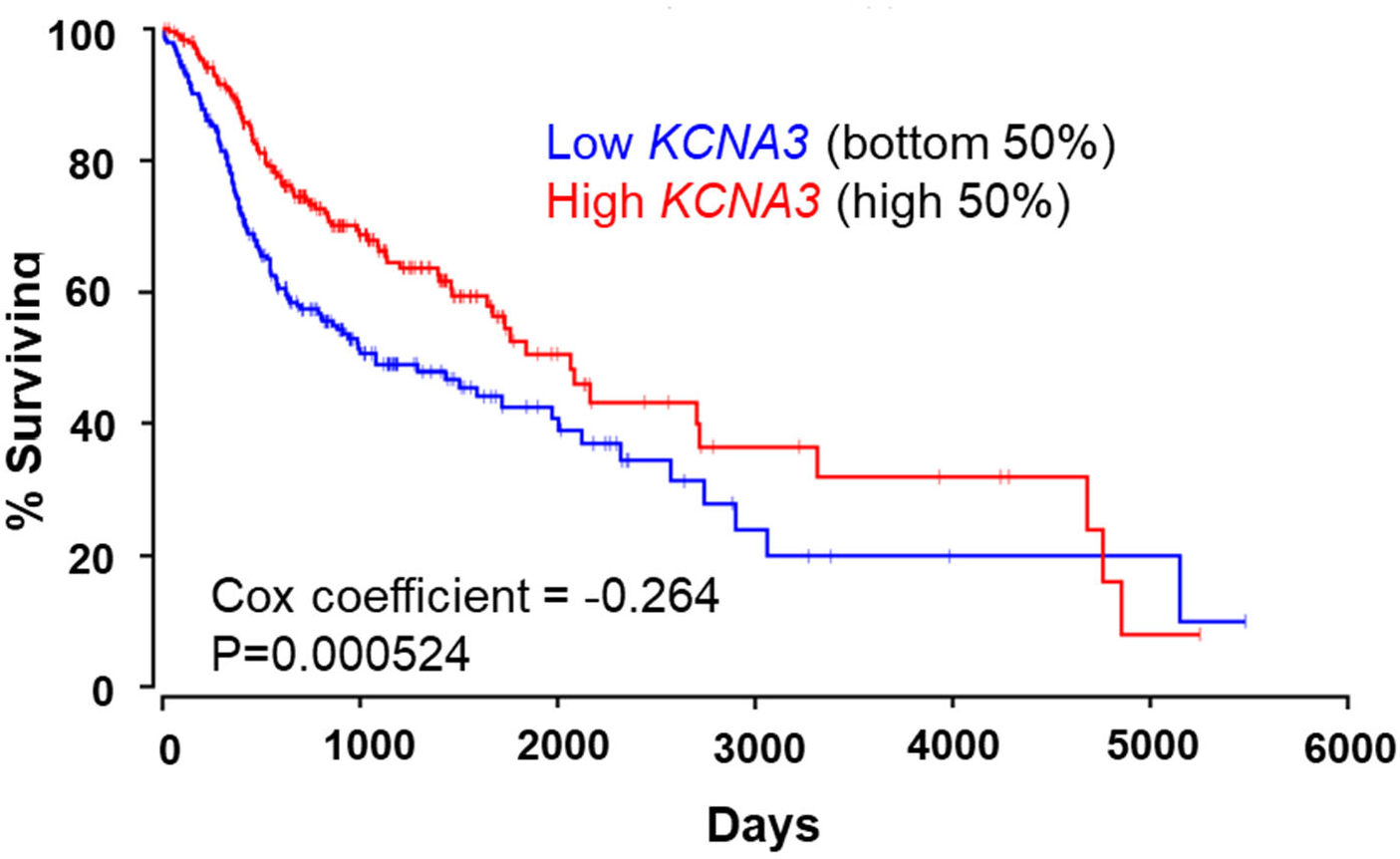
Association of KCNA3 expression with overall survival in TCGA HNSCC. Kaplan-Meier analysis of overall survival was performed on the TCGA HNSCC cohort, stratified by tumor KCNA3 expression. Patients were divided into KCNA3-low (bottom 50%) and KCNA3-high (top 50%) groups. Higher KCNA3 expression was significantly associated with improved overall survival compared to lower expression. Cox proportional hazards analysis revealed a negative coefficient (–0.264), indicating a protective effect of KCNA3 expression on survival (P = 0.000524).

### Design and fabrication of KCNA3-LNPs for targeted T cell therapy

A schematic overview of the general strategy for generating KCNA3-LNPs and testing their efficacy is shown in Fig. 2A. KCNA3-LNPs consist of a lipid shell and KCNA3 mRNA, which encodes the Kv1.3 potassium channel (Fig. 2B). The LNP shell is comprised of the ionizable cationic lipid 1,2-dioleoyl-3-trimethylammonium-propane chloride salt (DOTAP) for RNA encapsulation; l-α-phosphatidylcholine (PC), to facilitate lysosomal escape; PEGylated and biotinylated 1,2-distearoyl-sn-glycero-3-phosphoethanolamine-N-[biotinyl(polyethylene glycol)-2000] (PE-PEG-biotin) for in vivo stability (it decreases recognition by the mononuclear phagocytic system and clearance by renal filtration) and binding of targeting antibodies; cholesterol, as a stabilizer. The lipids were mixed at a molar ratio of 5:2:1:1 (DOTAP/PC/PE-PEG-biotin/cholesterol), the solution was then sonicated, and the resulting multilamellar vesicles were extruded through a 100 nm filter, as previously described (*21–23*). The LNP fabrication process is summarized in Fig. S2A. The LNPs were then functionalized with biotinylated anti-CD5 antibodies (isotype-matched IgG as a control, Fig. S2B). CD5 is expressed by T cells and a subset of B cells and anti-CD5 antibodies have been used to generate Chimeric Antigen Receptor (CAR) T cells in vivo (*24–26*). Dynamic light scattering using a Zetasizer Nano ZS was used to measure the LNP size and polydispersity index (PDI, the width of the particle distribution). The nanoparticle formulation exhibited a narrow and consistent size distribution (107 ± 12 nm, n=12 preparations) and low PDI (0.20 ± 0.05, n=9 preparations), indicating a high degree of sample homogeneity (Fig. S3). An average particle concentration of 7.19 ± 1.51 x 10^8^ particles/ml was determined for three independent reconstituted LNP formulations using nanoparticle tracking analysis (NTA) with NanoSight NS300.

**Fig. 2.**
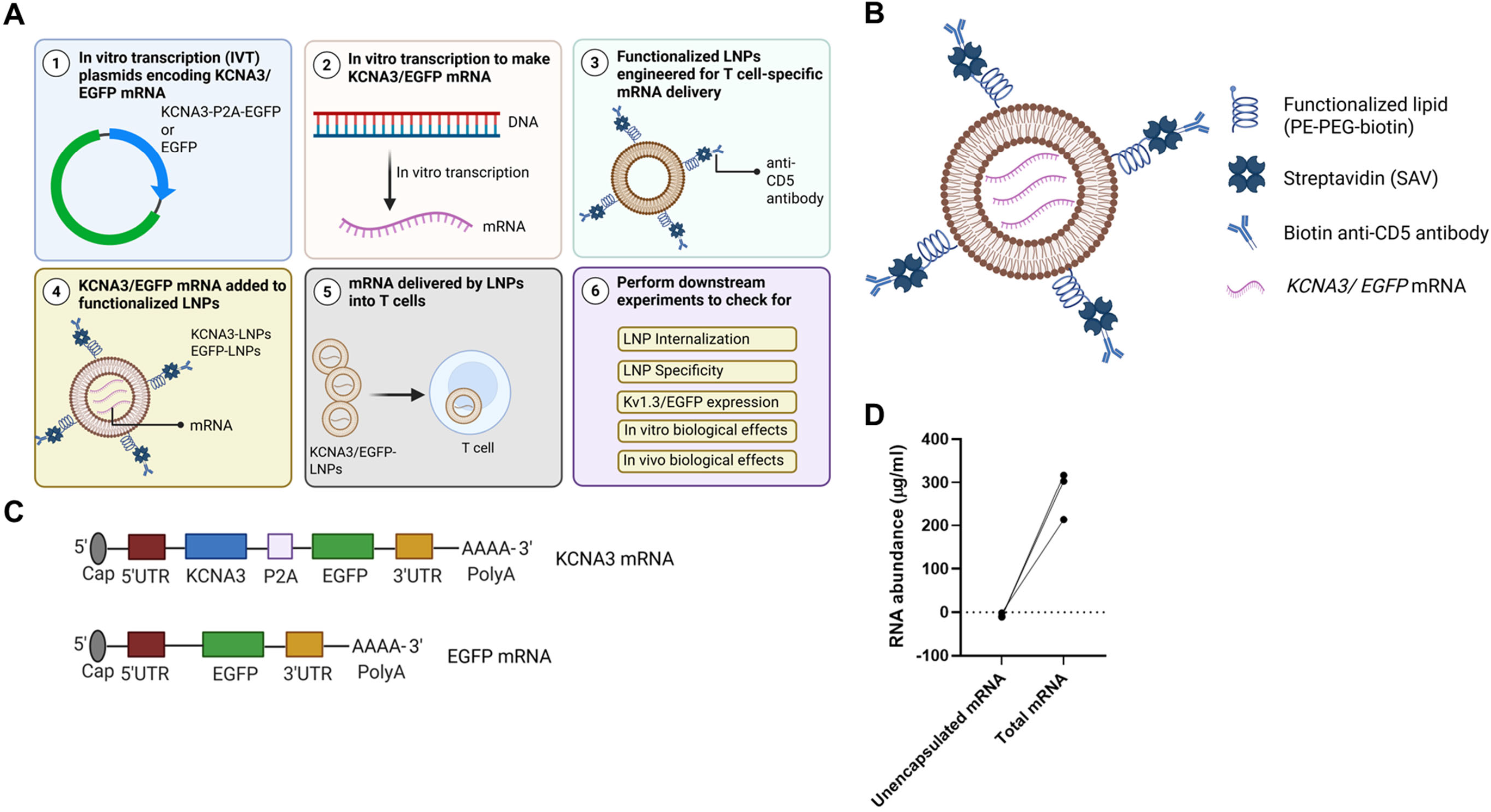
Design and characterization of KCNA3-LNPs and EGFP-LNPs. (A) A schematic representation for the strategy to generate KCNA3 or EGFP mRNA via IVT, followed by the engineering of functionalized LNPs for targeted delivery. This process involves synthesizing KCNA3 mRNA (mRNA encoding Kv1.3 channels), encapsulating it within LNPs designed for efficient transport into the desired cell population, and evaluating the efficacy of the delivery system in vitro and in vivo. (B) Schematic illustration of the LNP structure used to deliver KCNA3/EGFP mRNA (KCNA3-LNPs) or EGFP mRNA only (EGFP-LNPs; control). PE-PEG-biotin refers to 1,2-distearoyl-sn-glycero-3-phosphoethanolamine-N-[biotinyl(polyethylene glycol)-2000]. (C) Schematic representation of the in vitro–transcribed mRNA constructs that encode the Kv1.3 protein (the KCNA3 gene) and EGFP protein (EGFP gene). The KCNA3 mRNA includes a 5′ cap, 5′ untranslated region (UTR), KCNA3 coding sequence, a self-cleaving P2A linker, EGFP coding sequence, 3′ UTR, and a poly(A) tail. The EGFP control mRNA contains a 5′ cap, 5′ UTR, EGFP coding sequence, 3′ UTR, and a poly(A) tail. (D) Encapsulation of KCNA3 and EGFP mRNA within LNPs was assessed using a RiboGreen RNA quantification assay. RiboGreen fluorescence, measured in intact LNP preparations without Triton X-100, represents unencapsulated (free) mRNA, whereas samples treated with Triton X-100 to disrupt LNP membranes report total mRNA (encapsulated + free). Minimal RiboGreen signal was detected in the absence of detergent, while robust fluorescence was observed following Triton X-100 treatment, indicating that mRNA is predominantly encapsulated within LNPs. Data are representative of typical experiments performed independently in triplicate.

The KCNA3-LNPs contained an mRNA encoding both Kv1.3 and Enhanced Green Fluorescent Protein (EGFP), with EGFP serving as a marker for intracellular delivery (Fig. 2C). The DNA template for in vitro transcription (IVT) was engineered to include the coding sequences of KCNA3 and EGFP separated by a P2A sequence encoding a self-cleaving peptide to enable ribosomal skipping. Translation of this construct results in the co-expression of two distinct proteins: Kv1.3 and EGFP. EGFP-only mRNA containing LNPs (EGFP-LNPs) were used as controls. DNA constructs were synthesized in an IVT template plasmid containing a T7 promoter, 5′ and 3′ untranslated regions (UTRs), and a poly(A) tail for mRNA production. KCNA3 mRNA and control EGFP mRNA were obtained through IVT reaction followed by capping of the resulting mRNA. The workflow of the IVT reaction and capping is shown in Fig. S4 and is detailed in the Materials and Methods section. Prior to experimental application, the quality of mRNA obtained after IVT and capping was evaluated (Fig. S5), and only samples meeting the established quality control (QC) criteria (see Materials and Methods) were used in subsequent assays. This stringent QC guaranteed the integrity and consistency of the mRNA preparations, thereby ensuring that the experimental outcomes reported in the manuscript accurately reflect the effects of high-quality mRNA. The resulting mRNA (0.25 mg/ml) was incorporated into functionalized LNPs by rehydrating lyophilized particles with a solution containing mRNA and protamine, as previously described, and used for downstream experiments (*21*). mRNA encapsulation efficiency within the LNPs, quantified by the RiboGreen assay, was consistently >95% under these formulation conditions (Fig. 2D).

### Cell specificity and intracellular mRNA delivery of CD5 antibody-functionalized LNPs

To evaluate cell specificity, we compared LNPs labeled with biotinylated anti-CD5 antibodies (CD5-LNPs) and those labeled with biotinylated isotype control IgG antibodies (IgG-LNPs). These experiments utilized empty LNPs (LNPs lacking mRNA cargo), which were rendered fluorescent by conjugation with streptavidin Alexa Fluor 647 antibodies. Peripheral blood mononuclear cells (PBMCs) were isolated from the blood of healthy donors (HDs) and treated with fluorescent CD5-LNPs for 24 h (Fig. 3A) as previously described (*21–23*). Flow cytometry analysis revealed that CD5-LNPs selectively associated with CD4^+^/CD8^+^ T cell subsets and a small fraction of CD19^+^ B cells, but showed no interaction with CD16^-^CD14^+^ monocytes or CD56^+^ natural killer (NK) cells (Fig. 3B-C) (*27*). The IgG-LNPs did not bind to any of the immune cell subsets (Fig. 3D). The gating strategy used in these experiments is illustrated in Fig. S6.

**Fig. 3.**
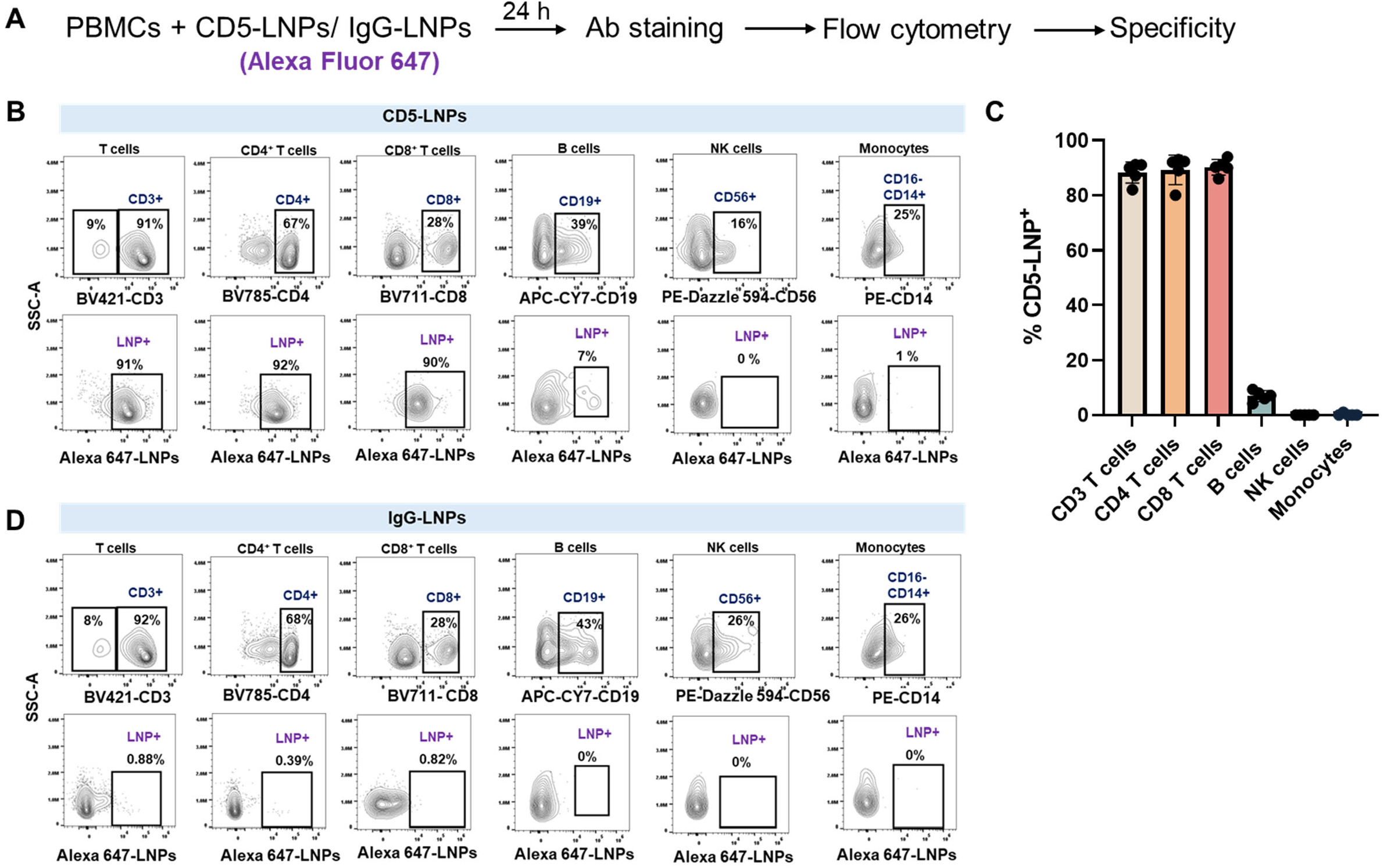
Selective association of CD5-coated LNPs with T cells. (A) The experimental protocol for measuring LNP specificity utilized Alexa Fluor 647-labeled LNPs functionalized with an anti-CD5 antibody (CD5-LNPs); isotype control biotin-conjugated mouse IgG1 antibody-functionalized LNPs served as controls (IgG-LNPs). PBMCs from HDs were incubated for 24 h with either CD5-LNPs or IgG-LNPs and stained the following day with cell-specific antibodies to identify immune cell subsets, as detailed in the Materials and Methods. LNP association within each immune cell subset was determined using flow cytometry. (B, D) Representative flow cytometry plots illustrate LNP^+^ populations expressing Alexa Fluor 647 fluorescence, achieved using Alexa Fluor 647 streptavidin (SAV) for LNP functionalization, in live CD3^+^, CD4^+^, CD8^+^ T cells, B cells (CD19^+^), NK cells (CD56^+^), and monocytes (CD16^-^CD14^+^) from PBMCs incubated for 24 h with CD5-LNPs (B) or IgG-LNPs (D). The gating strategy for these experiments to identify the various immune subsets is outlined in Fig. S6. Percentages represent the proportion of cells within each parent gate. (C) Quantification of CD5-LNP^+^ cells (measured by Alexa Fluor 647 fluorescence) in immune cell subsets across five HD PBMCs. Bar graphs represent the mean ± SD.

To assess intracellular mRNA delivery, we used confocal microscopy and flow cytometry, employing fluorescent EGFP-LNPs labeled with Alexa Fluor 647 (Fig. 4A). Confocal imaging of CD3^+^ T cells from HDs demonstrated EGFP protein expression in PBMCs treated with KCNA3- and EGFP-LNPs, but not in those treated with empty LNPs and no LNPs (Fig. 4B). These findings were corroborated by flow cytometry analysis which showed EGFP mRNA delivery by EGFP-LNPs in 87% of CD3^+^ T cells (Fig. 4C).

**Fig. 4.**
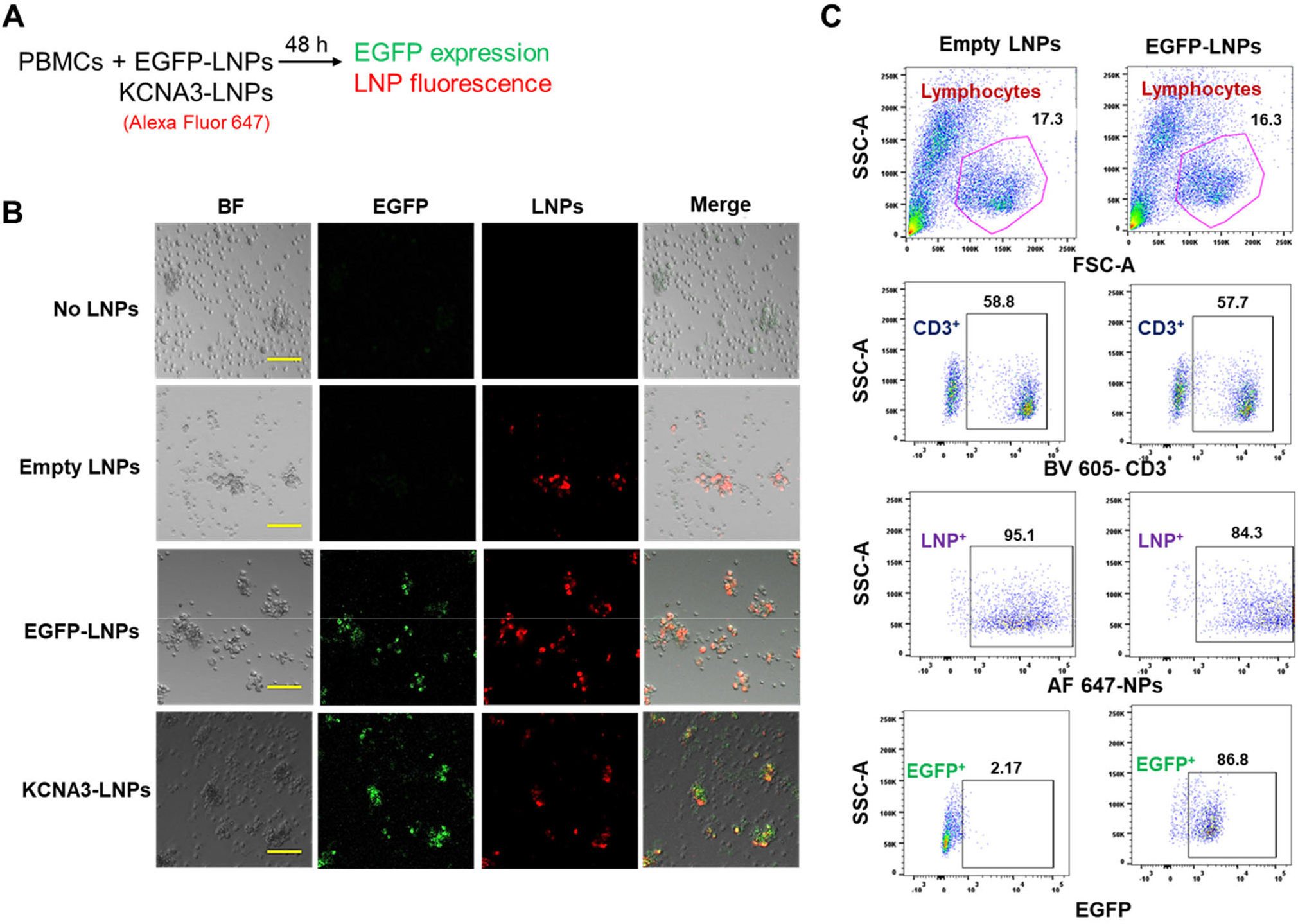
Selective delivery of LNP mRNA to T cells. (A) Experimental protocol for measuring intracellular delivery of Alexa Fluor 647-labeled LNPs functionalized with anti-CD5 antibodies and loaded with either EGFP mRNA (EGFP-LNPs) or mRNA encoding Kv1.3 channels and EGFP (KCNA3-LNPs), is illustrated. PBMCs from HDs were incubated for 48 h with EGFP-LNPs or KCNA3-LNPs, as detailed in the Materials and Methods section, and mRNA internalization was assessed via microscopy (B) or flow cytometry (C). HD PBMCs incubated with empty LNPs and PBMCs not incubated with any LNPs (no LNPs) served as controls. (B) Representative confocal images of HD PBMCs incubated with KCNA3-LNPs, EGFP-LNPs, empty LNPs, or no LNPs are shown, demonstrating EGFP expression (green) and LNP localization (red). Scale bar = 100 μm. (C) Representative flow cytometry histograms from a representative experiment showing EGFP expression in HD PBMCs treated with either EGFP-LNPs or Empty LNPs. The LNP^+^ population was identified by gating on the CD3^+^ T cell population expressing NP fluorescence (Alexa Fluor 647). The numbers above each gate represent the percentage of cells within the gate. The experiments were conducted independently using PBMCs from three HD PBMCs. Data shown are representative of one independent experiment.

### Increased functional Kv1.3 channels in CD3⁺ T Cells following KCNA3-LNP exposure

Flow cytometry confirmed the efficient delivery of mRNA encoding Kv1.3 channels to CD3^+^ T cells by KCNA3-LNPs (Fig. 5A). PBMCs from HDs were treated with fluorescently labeled KCNA3-LNPs, EGFP-LNPs, or empty LNPs, or left untreated (no LNPs). Cells were treated with LNPs at a final concentration of 1.20 x 10^8^ particles/ml, and a cell-to-particle ratio of 1:180, along with 41.67 μg/ml encapsulated mRNA for KCNA3- and EGFP-LNPs (see Materials and Methods for calculations). The experimental protocol and the flow cytometry gating strategy used in these experiments are shown in Fig. S7A and B, respectively. Kv1.3 protein abundance increased more than two-fold in KCNA3-LNP-treated T cells, whereas the geometric mean fluorescence intensity (gMFI) in EGFP-LNP- and empty LNP-treated cells was comparable to that of untreated controls (Fig. 5A). This increase in Kv1.3 protein was not accompanied by changes in the early activation marker CD69 (Fig. 5B).

**Fig. 5.**
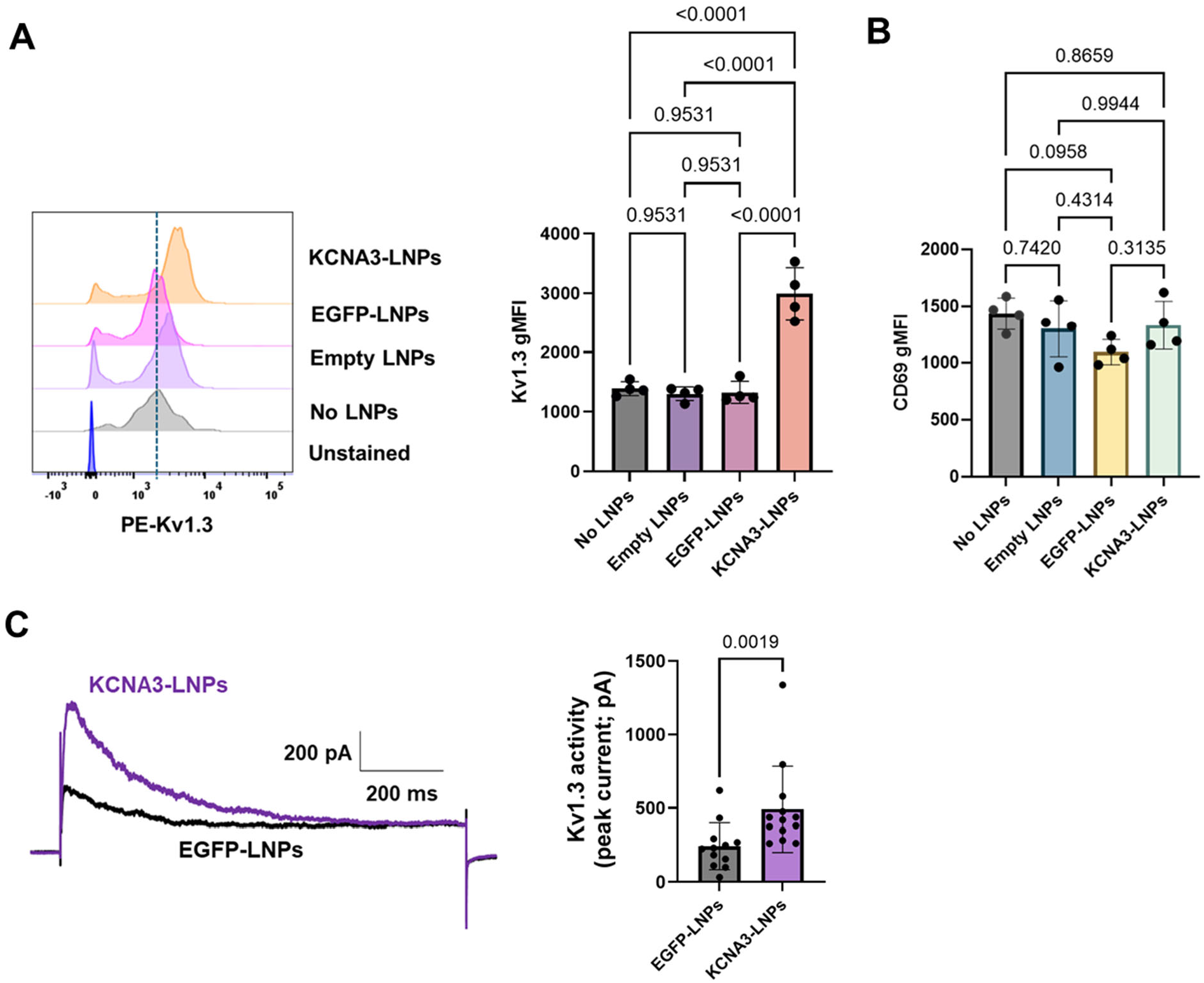
Increased Kv1.3 expression and activity in T cells by KCNA3-LNPs. PBMCs from HDs were incubated for 48 h with either EGFP-LNPs or KCNA3-LNPs (1.20 x 10^8^ particles/ml; 41.67 μg/ml encapsulated mRNA) as detailed in Materials and Methods. We then measured Kv1.3 abundance (using flow cytometry) and activity (using electrophysiology), along with CD69 abundance (using flow cytometry), within the CD3^+^ population. PBMCs treated with LNPs without mRNA (empty LNPs) and PBMCs not treated with any LNPs (no LNPs) served as experimental controls. The flow cytometry experimental protocol and gating strategy are outlined in Fig. S7. (A) (Left) Staggered histograms from a representative experiment demonstrate Kv1.3 expression in the CD3^+^ T cell population of PBMCs treated with KCNA3-LNPs, EGFP-LNPs, empty LNPs, or no LNPs. (Right) The average geometric mean fluorescence intensity (gMFI) of surface Kv1.3 expression in CD3^+^ T cells from four HDs treated with KCNA3-LNPs, EGFP-LNPs, empty LNPs, or no LNPs was determined. Significance was assessed using one-way ANOVA (P<0.0001). Post-hoc testing was performed using the Holm-Sidak multiple comparison test. (B) The average gMFI of CD69 expression in CD3^+^ T cells from four HDs treated with KCNA3-LNPs, EGFP-LNPs, empty LNPs, or no LNPs was determined. Significance was assessed using one-way ANOVA (P= 0.1233). Post-hoc testing was performed using Tukey’s test. (C) Increased Kv1.3 channel activity in CD3^+^ T cells by KCNA3-LNPs. Representative Kv1.3 currents elicited by step pulse depolarization to +40 mV from -80 mV holding potential (HP) in CD3^+^ T cells from the same donor are shown on the left, and peak Kv1.3 currents measured in CD3^+^ T cells treated with KCNA3-LNPs (n=13 cells) and EGFP-LNPs (n=12 cells) from two HDs are shown on the right. Significance was determined using the Mann-Whitney U test. Bar graphs represent the mean ± SD.

To determine whether the increase in Kv1.3 protein expression in KCNA3-LNP treated cells corresponded to an increase in functional Kv1.3 channels, we measured the electrical activity of these channels using patch-clamp electrophysiology (Fig. 5C). Kv1.3 channels open upon membrane depolarization and undergo a rapid current decline during sustained depolarization, a process termed C-type inactivation (*28*). Currents recorded from KCNA3-LNP treated cells displayed the characteristic features of wild-type Kv1.3 channels (Fig. 5C left). Importantly, current amplitude was doubled in KCNA3-LNP treated T cells compared with that in EGFP-LNP controls (Fig. 5C right). Cell capacitance, a measure of cell size and an indicator of T cell activation (*13*), was modestly increased in KCNA3-LNP treated cells (2.2 ± 0.2 pF, n=13 cells from two HDs, mean ± SEM) compared with EGFP-LNP treated controls (1.7 ± 0.1 pF, n=12 cells from two HDs; P=0.049, mean ± SEM, Welch’s t-test). Despite this, Kv1.3 current density remained significantly higher in KCNA3-LNP treated cells (235.8 ± 33 pA/pF, n=13 cells from two HDs, mean ± SEM) relative to controls (153.5 ± 32.3 pA/pF; n=12 cells from two HDs; P=0.0298, mean ± SEM, Mann-Whitney U test).

### Enhanced Ca^2+^ signaling and cytokine and cytotoxic molecule release in CD8^+^ T cells by KCNA3-LNPs

An increase in Kv1.3 channels is known to enhance T cell functional capabilities by promoting greater Ca^2+^ influx (*5, 6, 29*). Flow cytometry experiments using the Ca^2+^ indicator Indo-1 demonstrated that KCNA3-LNPs (1.20 x 10^8^ particles/ml; 41.67 μg/ml encapsulated mRNA) induced a 50% increase in Ca^2+^ influx in HD CD8^+^ T cells (Fig. 6A). Ca^2+^ influx was triggered using thapsigargin (TG) in Ca^2+^-free medium, followed by the reintroduction of extracellular Ca^2+^. TG, an inhibitor of the sarco/endoplasmic reticulum (ER) Ca^2+^-ATPase (SERCA) pump, depletes ER Ca^2+^ stores and consequently activates Ca^2+^-release activated Ca^2+^ (CRAC) channels, allowing Ca^2+^ influx into the cell upon reintroduction of Ca^2+^ in the extracellular milieu (*5, 11*). This technique is commonly used to assess the ability of T cells to flux Ca^2+^ through ion channels in a T cell receptor (TCR) -independent manner (*13, 14, 21*).

**Fig. 6.**
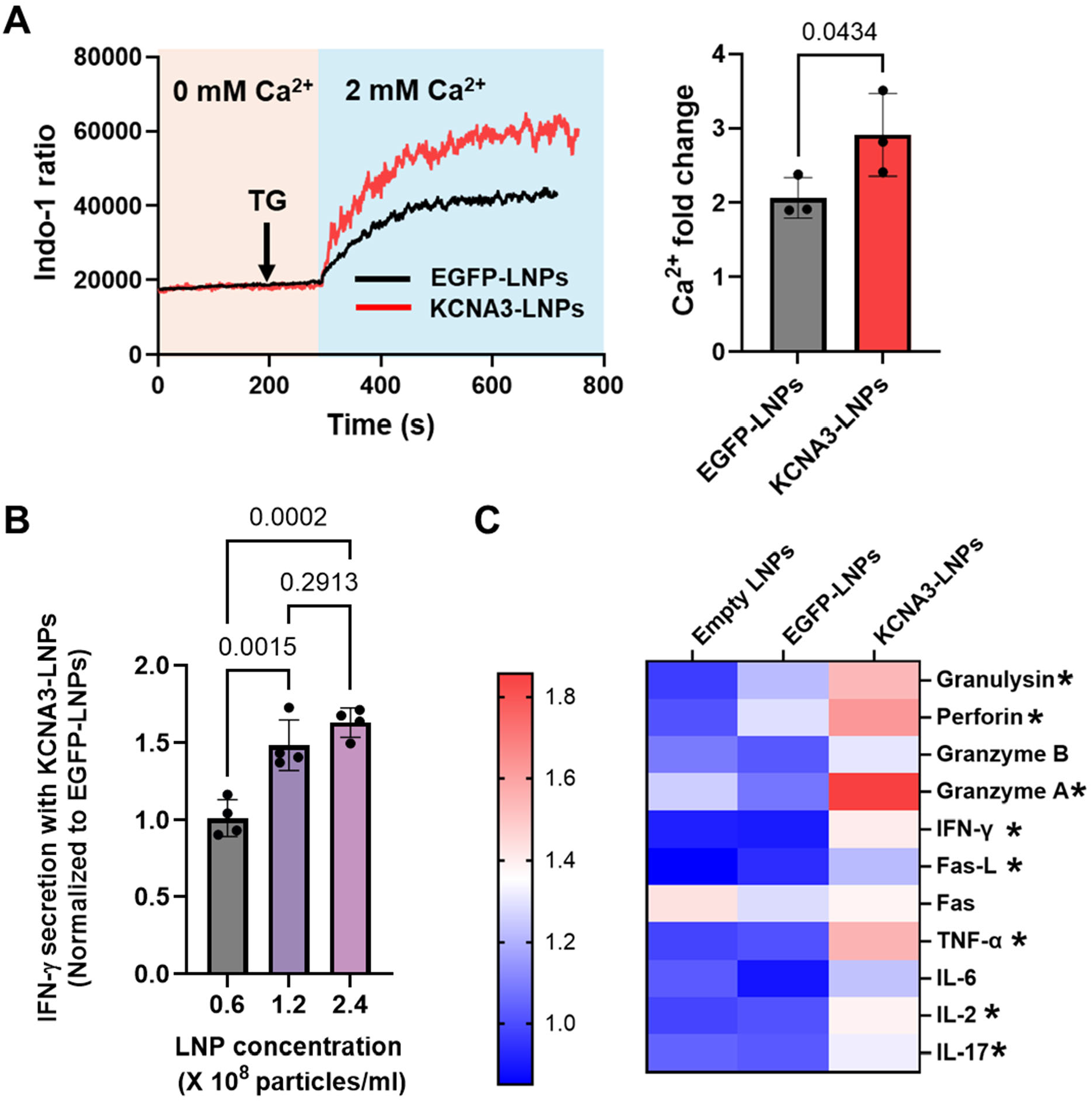
Enhanced Ca^2+^ response and cytokine production in T cells by KCNA3-LNPs. (A) The left panel shows a representative Ca^2+^ response, reported as a ratio of Indo-1 fluorescence at 400 and 480 nm, recorded in activated CD3^+^ T cells from HDs treated with either KCNA3-LNPs or EGFP-LNPs (1.20 x 10^8^ particles/ml; 41.67 μg/ml encapsulated mRNA) for 48 h. Cells were loaded with the Indo-1 ratiometric dye, and fluorescence was measured by flow cytometry. Initially, Indo-1 loaded cells were exposed to thapsigargin (TG, indicated by an arrow) in 0 mM Ca^2+^, followed by 2 mM Ca^2+^, resulting in a rapid Ca^2+^ influx (refer to Materials and Methods). The right panel displays the average fold change in peak Ca^2+^ levels. The data represent independent experiments conducted on CD3^+^ T cells isolated from n=3 HDs. Significance was assessed using an unpaired t-test. (B) IFN-γ production by HD CD8⁺ T cells treated with increasing concentrations of KCNA3-LNPs or EGFP-LNPs (0.6 x 10^8^, 1.20 x 10^8^ and 2.4 x 10^8^ particles/ml, and 20.83 μg/ml, 41.67 μg/ml and 83.34 μg/ml encapsulated mRNA). Cells were then activated for 48 h. IFN-γ levels were quantified by ELISA and expressed as fold change relative to the average IFN-γ concentration in EGFP-LNP–treated controls. Data represent mean ± SD from n = 4 HDs. Statistical significance was determined by one-way ANOVA (P = 0.0002) with Tukey’s multiple-comparisons test. (C) CD8^+^ T cells from eight HDs were activated for 48 h and treated with KCNA3-LNPs, EGFP-LNPs (1.20 x 10^8^ particles/ml; 41.67 μg/ml encapsulated mRNA), empty LNPs (1.20 x 10^8^ particles/ml), or no LNPs. Cytokines and cytotoxicity biomarkers were measured in the supernatants using the Human CD8/NK LEGENDplex flow cytometry-based multiplex immunoassay. The heatmap illustrates the fold change in analyte concentration in the KCNA3-LNPs, EGFP-LNPs, and empty LNP-treated groups compared to the no LNP-treated group. Data were obtained from eight HDs, with each donor receiving all LNP treatments. Statistical significance was determined using one-way analysis of variance (ANOVA) or Kruskal-Wallis test, where samples did not meet normality. Post-hoc testing was conducted using Dunn’s test, and statistical significance between the KCNA3-LNPs- and EGFP-LNPs-treated groups is indicated by an asterisk (*).

Ca^2+^ influx is one of the earliest events following TCR stimulation and plays a key role in regulating gene expression, including cytokines and exocytosis (*11, 14, 30*). KCNA3-LNPs resulted in an increase in IFN-γ secretion by activated human CD8⁺ T cells compared with control LNPs (Fig. S8), suggesting their potential to enhance cytotoxic T cell function. In contrast, treatment of cells with no LNPs, empty LNPs, or EGFP-LNPs did not affect IFN-γ release (Fig. S8). The concentration of KCNA3-LNPs used in the Ca^2+^ flux experiments elicited a maximal increase in IFN-γ release by T cells (Fig. 6B) and was therefore applied in all subsequent assays. ELISA measurements reflected IFN-γ secretion from the entire cell population, including ∼10% of the cells that did not internalize KCNA3-LNPs (Fig. 3B). Consequently, the observed 50% increase in IFN-γ likely underestimated the true effect of KCNA3-LNPs on successfully transfected cells.

Multiplex immunoassay analysis showed that treatment with KCNA3-LNPs (1.20 x 10^8^ particles/ml; 41.67 μg/ml encapsulated mRNA), significantly enhanced the release of effector cytokines (IL-2, IL-17, TNF-α, and IFN-γ), Fas ligand (FasL), and cytotoxic molecules (Granzyme A, perforin, and granulysin) from activated human CD8⁺ T cells (Fig. 6C). These results provide in vitro evidence supporting the antitumor efficacy of KCNA3-LNPs, as effector molecules upregulated by KCNA3-LNPs are critical for tumor cell killing (22).

### Effective antitumor response following local KCNA3-LNP delivery in tumor-bearing mice

In vivo experiments were conducted using humanized PBMC (hu-PBMC)-HNSCC xenograft mice to assess the antitumor efficacy of the KCNA3-LNPs. Immunodeficient NOD/LtSz-SCID IL-2RG−/− (NSG) mice were flank-injected with tumor spheroids derived from immortalized human Cal27 HNSCC cells (Fig. 7A). Upon tumor formation, HD PBMCs were engrafted and, after 24 h, KCNA3- or EGFP-LNPs were injected peritumorally every 3-4 days (7.19 x 10^8^ particles/ml; 100 μl) (Fig. 7A). Tumor development was monitored for 17 days after the first LNP injection (or for 41 days after spheroid injection), after which the mice were sacrificed, and tumors were removed and stained for pathology and T cell function assessments (Fig. 7A).

**Figure 7.**
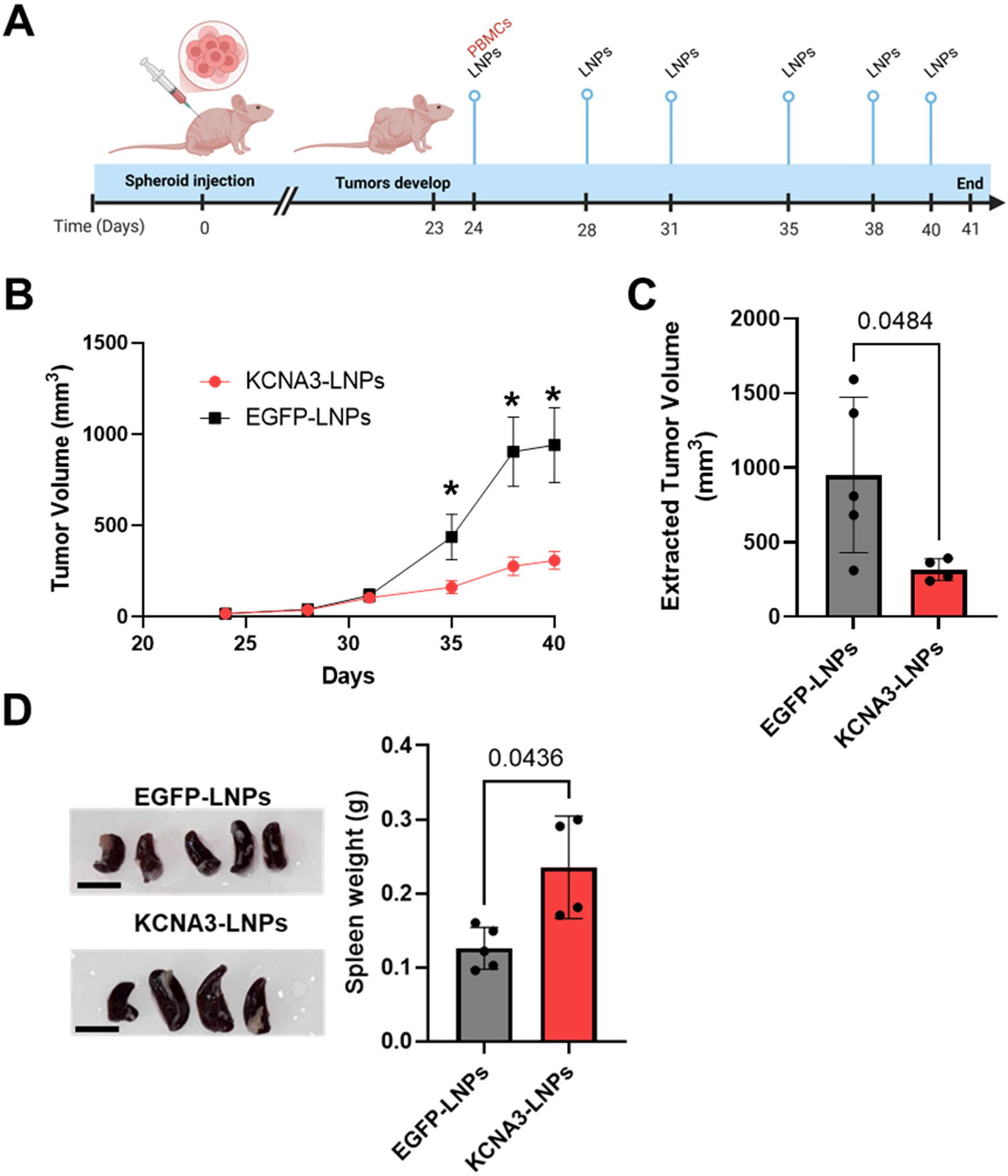
Reduction in tumor burden by KCNA3-LNP treatment in a hu-PBMC-HNSCC xenograft mouse model. (A) A schematic representation of the hu-PBMC-HNSCC mouse studies is presented. NSG mice were injected with Cal 27 spheroids, resulting in palpable tumors by day 23 post-injection. Subsequently, the mice were engrafted with HD PBMCs via the tail vein and peritumoral injections; KCNA3-LNPs or control EGFP-LNPs were then administered peritumorally (7.19 x 10^8^ particles/ml; 100 μl) every 3–4 days, on the days indicated in the schematic. Tumor volume was measured on treatment days until the study endpoint (day 41). (B) Line graphs illustrate the tumor volumes measured throughout the course of the study in mice injected with KCNA3-LNPs (n=4) and EGFP-LNPs (n=5). Data are presented as mean ± SEM, and statistical significance between groups was determined by Welch’s t-test, indicated by an asterisk (*). (C) At the study endpoint on day 41, tumors were excised, and their volumes were measured as described in the Materials and Methods. The extracted tumor volumes from mice injected with EGFP-LNPs (n=5) and KCNA3-LNPs (n=4) are displayed in a bar graph, with data presented as means ± SD. Statistical significance was assessed using an unpaired t-test. (D) Images on the left show spleens dissected from mice injected with EGFP-LNPs (n=5) and KCNA3-LNPs (n=4), with a scale bar of 10 mm. The bar graph on the right illustrates the weights of the dissected spleens from EGFP-LNP- and KCNA3-LNP-injected mice, with data shown as means ± SD. Statistical significance was determined using an unpaired t-test.

Tumor burden was decreased in KCNA3-LNP treated mice as compared to that in EGFP-LNP treated mice (Fig. 7B) and the volume of the tumors extracted at the end of the experiment (Day 41) was also significantly lower in KCNA3-LNP treated group (Fig. 7C). There was no difference in tumor volumes between EGFP-LNP mice and mice that were not injected with LNPs (Fig. S9). Furthermore, there was a significant increase in spleen weight in the KCNA3-LNP treated group (Fig. 7D), but not in the other organs (Fig. S10).

Next, we evaluated the impact of KCNA3-LNP treatment on antitumor immunity in vivo in hu-PBMC-HNSCC mice (same treatment as in Fig 7A). Tumors harvested at the study endpoint were stained for histopathological and immunohistochemical analyses (Fig. 8A). Hematoxylin and eosin (H&E) staining of formalin-fixed paraffin-embedded (FFPE) tumor sections revealed significant differences in the degree of peritumoral inflammatory response. Tumors from KCNA3-LNP-treated mice displayed prominent, thick bands of peritumoral inflammatory infiltrates compared with those from EGFP-LNP-treated controls which more often showed sparse, thin bands of peritumoral inflammation (Fig. 8B and Fig. S11). Immune cell infiltrates showed no differences between EGFP-LNP-injected mice and mice that were not injected with LNPs (Fig. S12). Semi-quantitative histopathological scoring (Table S1) further demonstrated that although inflammatory cell infiltrates were detectable across all groups, KCNA3-LNP treatment resulted in a notably higher accumulation of tumor-infiltrating lymphocytes localized to the tumor periphery, consistent with an enhanced immune-engaged TME (Fig. 8C). Immunohistochemical analysis of tumor sections showed a significant increase in intratumoral CD8⁺ T cell abundance in KCNA3-LNP–treated mice compared to EGFP-LNP-treated controls (Fig. 8D, E). Notably, a substantial proportion of these infiltrating CD8⁺ T cells in KCNA3-LNP-treated tumors also co-expressed Granzyme B, indicating an enhanced cytotoxic phenotype and functional activation within the tumor (Fig. 8D, F). In contrast, EGFP-LNP-control tumors and tumors from mice not injected with LNPs exhibited fewer CD8⁺ T cells with limited Granzyme B expression, consistent with a less effective antitumor immune response (Fig. 8D-F).

**Fig. 8.**
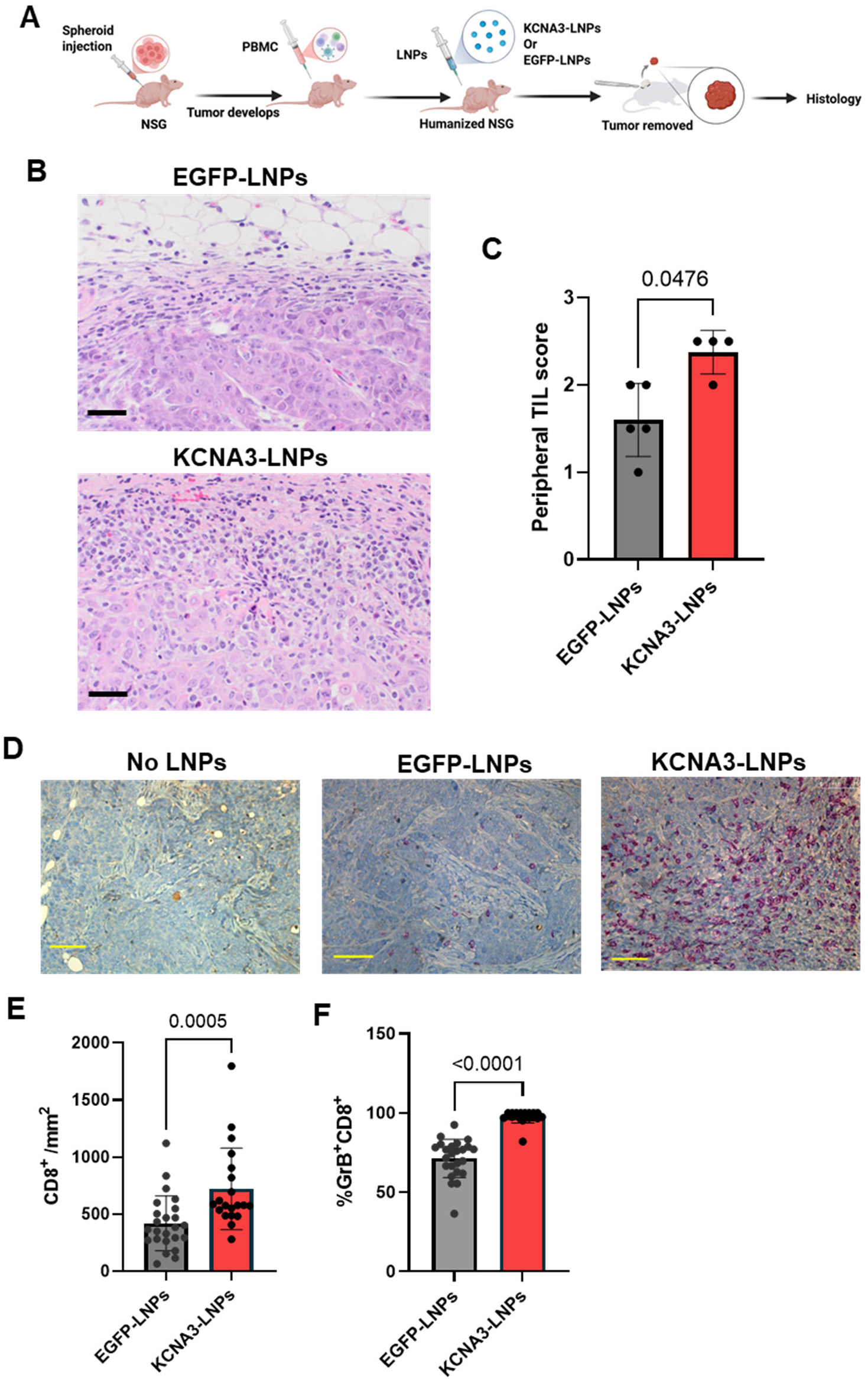
Enhanced intratumoral presence of functionally competent cytotoxic T cells in a hu-PBMC-HNSCC xenograft mouse model by KCNA3-LNPs. **(A) A schematic** representation of the hu-PBMC-HNSCC mouse studies is shown. NSG mice were injected with Cal 27 spheroids, leading to the development of palpable tumors. Subsequently, the mice were engrafted with HD PBMCs via tail vein and peritumoral injections and then received peritumoral injections of KCNA3-LNPs or control EGFP-LNPs (7.19 x 10^8^ particles/ml; 100 μl) every 3–4 days. Tumors were excised at the study endpoint and processed for histological analysis. (B) Representative hematoxylin and eosin-stained tumor tissue from hu-PBMC-HNSCC mice treated with EGFP-LNPs or KCNA3-LNPs demonstrates increased inflammatory cell abundance within the tumor in the latter. Detailed annotations of these images are provided in Fig. S11. Scale bar = 50 μm. (C) The abundance of tumor infiltrating lymphocytes (TILs) in the periphery of the tumors was scored by a board-certified pathologist (see Materials and Methods and Table S1) in KCNA3-(n= 4) and EGFP-LNPs-treated (n= 5) mice. The bars represent the mean ± SD, and significance was determined by the Mann-Whitney U test. (D) IHC of the tumor tissue stained for CD8 (brown signal) and Granzyme B (red signal) in representative tumor sections from mice treated with KCNA3-LNPs, EGFP-LNPs, and untreated (no LNPs) mice. Scale bar = 50 μm. (E) Intratumoral abundance of CD8^+^ T cells in KCNA3-LNPs (n= 4) and EGFP-LNPs-treated (n= 5) mice, calculated as the number of cells divided by the area of the regions of interest (ROIs) within the intratumoral region of the tissues. (F) Granzyme B^+^CD8^+^ T cell abundance in KCNA3-LNPs (n= 4) and EGFP-LNPs-treated (n= 5) mice, normalized for the area of ROIs. For each biopsy, five fields (ROIs) were imaged, and significance was determined by unpaired Student’s t-test. The bars represent the mean ± SD, and each symbol represents an individual measurement from the analyzed fields.

These data collectively demonstrate that local delivery of KCNA3-LNPs transforms the tumor immune microenvironment by enhancing the recruitment of cytotoxic CD8⁺ T cells. This process ultimately leads to improved tumor-immune engagement and is associated with a reduced tumor burden. These findings establish LNP-based KCNA3 mRNA delivery as a powerful strategy to restore antitumor T cell function in an immunosuppressive solid tumor setting, providing both in vitro and in vivo validation for targeting Kv1.3 potassium channel-dependent immune dysfunction as a potential therapeutic modality for HNSCC.

## DISCUSSION

Immunotherapy has transformed cancer treatment; however, durable responses in solid tumors, such as HNSCC, remain limited, with most patients not benefiting from conventional immunotherapy alone (*4*). A major barrier is the immunosuppressive TME, which impairs T cell Ca^2+^ signaling, cytotoxicity, cytokine secretion, and migration (*13, 14, 29*). Here, targeted delivery of KCNA3 mRNA via LNPs selectively enhanced Kv1.3 channel expression in T cells, boosting Ca^2+^ signaling, a critical early step in activation (*5, 11*). Consequently, the secretion of key cytokines and cytotoxic molecules, a Ca^2+^-dependent process was increased, leading to improved T cell antitumor functions (*11, 14, 30*). Localized peritumoral KCNA3-LNP administration in a preclinical HNSCC model reduced tumor burden, restored robust T cell–mediated antitumor immunity, and increased intratumoral Granzyme B⁺ CD8⁺ T cell infiltration, demonstrating that KCNA3-LNPs reprogram the TME toward an antitumor phenotype.

Kv1.3 potassium channels are critical regulators of T cell survival, proliferation, and effector function, including cytokine production and cytotoxicity, particularly in effector memory T cells (*5, 6*). Although expressed in regulatory T cells, their contribution to suppressive function is minimal (*31*). In HNSCC, TME-mediated downregulation of Kv1.3 in cytotoxic T cells limits antitumor immunity and response to anti-PD1 therapy (*13, 14*). Restoring Kv1.3 activity in T cells is therefore critical for re-establishing effective antitumor immunity. In contrast, Kv1.3 is overexpressed in many tumors, including HNSCC, supporting tumor cell proliferation and survival via plasma membrane and mitochondrial functions (*19, 32–34*). From the perspective of tumor cells, Kv1.3 inhibition is therefore predicted to exert a strong antitumor effect. This opposing role of Kv1.3 underscores the need for precise targeting, as indiscriminate elevation could enhance tumor survival or affect other cell populations expressing Kv1.3, including monocytes and platelets (*18, 35*). Our anti-CD5-functionalized LNPs restricted KCNA3 overexpression predominantly in T cells, minimizing off-target exposure (*25, 26*). Furthermore, transient mRNA expression may limit long-term adverse effects (*36, 37*).

LNP-based, cell-targeted mRNA delivery offers several advantages. Our biocompatible LNPs enable efficient cytosolic delivery of mRNA following cellular uptake. Composed of ionizable lipids, helper lipids, cholesterol, and PEGylated lipids, LNPs protect the cargo from degradation while enhancing cellular uptake and promoting endosomal escape, features that are central to the clinical success of mRNA vaccines and other nucleic acid–based therapeutics (*16*). We have previously utilized a similar LNP platform to deliver oligonucleotides targeting Kv1.3 in effector memory T cells, achieving selective channel silencing and therapeutic benefit in a murine lupus nephritis model (*23*). Building on this experience, we repurposed the platform for Kv1.3 augmentation, incorporating CD5-targeted functionalization to selectively direct mRNA delivery to T lymphocytes. Furthermore, KCNA3-LNPs were engineered to allow unstable mRNA to be loaded into lyophilized LNPs immediately before administration, thereby combining the potential for extended shelf life with minimized cargo degradation (*21, 38*). The particle concentration and mRNA incorporation were quantified in three independent formulations. The consistent observation of functional outcomes across various independent formulations underscores the reproducibility and stability of the platform. Pharmacostability studies are necessary to further validate this formulation strategy. In addition to molecular delivery, LNPs offer substantial translational advantages at the therapeutic level. Compared with adoptive T cell transfer, LNP delivery simplifies logistics, reduces costs, improves scalability, and provides transient, non-genetically permanent T cell reprogramming.

Anti-CD5 antibodies were selected as the cell-targeting moieties. CD5 is a surface glycoprotein expressed on T cells and a subset of B cells (*25, 26*). In T cells, CD5 negatively regulates TCR signaling, shaping thymic selection, maintaining self-tolerance, and limiting excessive activation (*25, 26*). This characteristic makes CD5 an attractive target for immune modulation, including engineered T cells and Treg strategies (*39–42*). Consistent with this, CD5-functionalized LNPs were preferentially associated with CD3⁺ T cells, with limited uptake by CD19⁺ B cells. Although CD5⁺ B cells may contribute to tumor-associated immune regulation, their role in human solid tumors remains unclear (*25, 26, 43*). However, in hu-PBMC-engrafted mice, B cells are present only at minimal levels and fail to persist over time, resulting in a human leukocyte population that is predominantly composed of T cells (*44–46*). Overall, the observed T cell selectivity supports CD5 as a safe and effective targeting handle, and EGFP-LNP controls showed no functional effects attributable to the targeting moiety itself.

Functionally, KCNA3-LNPs increased Ca^2+^ signaling and enhanced cell cytotoxicity and cytokine secretion by augmenting Kv1.3 in T cells. KCNA3-LNP treatment doubled the Kv1.3 current amplitude and increased channel protein expression without inducing early activation markers such as CD69, indicating functional potentiation without nonspecific early T cell activation, which can drive terminal differentiation (*13*). Overall, KCNA3-LNPs increased the levels of IL-2, IFN-γ, TNF-α, and IL-17, supporting their proliferation, survival, and antitumor activity. Concurrent upregulation of perforin, granzymes, granulysin, and the Fas ligand enhanced both granule-mediated and death-receptor cytotoxic pathways (*47*). Together, these changes demonstrate that KCNA3-LNPs reshape the cytokine milieu toward an antitumor phenotype.

The efficacy of KCNA3-LNP delivery is demonstrated by increased intratumoral Granzyme B⁺ CD8⁺ T cell infiltration and reduced tumor burden in an HNSCC mouse model. These effects were achieved through localized peritumoral administration, which was selected to maximize LNP delivery to tumor-draining lymph nodes and the TME. Although this approach, which is already utilized clinically in HNSCC, may limit off-target effects, it could restrict systemic efficacy in metastatic settings (*48, 49*). Nonetheless, the splenomegaly observed in the treated mice suggests systemic immune modulation. Although human PBMC engraftment in NSG mice can eventually lead to graft-versus-host disease (GvHD) (*50*), the increased spleen size in KCNA3-LNP-treated mice is consistent with expansion of functional human effector T cells rather than GvHD, as clinical signs of disease progression, including impaired mobility and cachexia, were absent. Splenomegaly was not observed in control mice. Splenic enlargement in KCNA3-LNP-treated mice correlated with higher intratumoral CD8⁺ density and Granzyme B expression, indicating that systemic T cell priming enhances local tumor infiltration. Future studies should optimize systemic efficacy, biodistribution, and target engagement and explore combinations with anti-PD1 or CAR-T therapies. Broad profiling of the TME and long-term safety studies are critical for clinical translation.

In conclusion, the clinical success of mRNA-LNP technology in COVID-19 vaccines provides a strong foundation for KCNA3-LNP translation (*16*). Modular and scalable LNP platforms facilitate rapid adaptation to diverse cargos and targets. Selective Kv1.3 augmentation represents a form of in vivo T cell engineering that enhances endogenous effector machinery rather than introducing synthetic receptors, potentially overcoming immunosuppression in solid tumors. This approach may be extended to other malignancies characterized by immunosuppressive TME, such as ccRCC, in which high KCNA3 expression correlates with improved survival. Overall, these proof-of-concept studies demonstrated the specificity, efficacy, and safety of KCNA3-LNPs and supported the further development of clinical trials to improve patient outcomes in solid tumors.

## MATERIALS AND METHODS

### Study Design

This study aimed to assess whether targeted delivery of KCNA3 mRNA (mRNA encoding Kv1.3 channels) via CD5-functionalized lipid nanoparticles (KCNA3-LNPs) could restore Kv1.3 channel function in T cells and enhance antitumor immunity in HNSCC. The experimental design incorporated both in vitro and in vivo methodologies. In vitro investigations utilized human PBMCs from HDs to evaluate nanoparticle specificity, intracellular mRNA delivery, Kv1.3 protein levels and channel activity, Ca^2+^ signaling, and subsequent cytokine and cytotoxic molecule production. Functional assessments included flow cytometry, electrophysiology, confocal microscopy, ELISA, multiplex cytokine assays, and Ca^2+^ flux measurements. For in vivo therapeutic efficacy evaluation, a humanized HNSCC mouse model was developed by engrafting immunodeficient NSG mice with Cal27 tumor spheroids, followed by transfer of human PBMCs (hu-PBMC-HNSCC mice). Subsequently, the mice received peritumoral injections of either KCNA3-LNPs or control EGFP-LNPs. Treatment effects were determined by evaluating the tumor burden, histopathology, immune cell infiltration, and markers of T cell cytotoxicity. Mice were randomly allocated to treatment groups. The sample sizes were determined based on pilot experiments and prior experience with this model, ensuring sufficient statistical power to detect statistically significant differences. No samples were excluded from the analysis, and blinding was not performed except for histopathologic scoring. Statistical analyses are detailed in the Statistical analysis section, with specific statistical tests and P values provided in the corresponding figures and figure legends.

### TCGA Analysis

TCGA analysis was performed using OncoLnc (www.oncolnc.org), accessed in March 2020. OncoLnc is an interactive tool that assesses correlations between clinical and expression data of mRNA, miRNAs, and lncRNAs from over 8500 patients across 21 TCGA cancer studies (*51*). KCNA3 was selected as the target gene. A Cox regression analysis was performed for KCNA3 across the 21 cancer types, with a negative Cox coefficient indicating a protective effect against the disease of interest. Subsequently, Kaplan-Meier curves were plotted for KCNA3 in HSNCC (Fig. 1) and ccRCC (Fig. S1). Patients were selected within the upper and lower quartiles of the included cohort, and Kaplan-Meier plots were generated for each percentile group, including the sample size (n) and log-rank P-value.

### IVT plasmid constructs

Plasmids encoding human KCNA3 (Kv1.3 protein) and EGFP for IVT were custom-designed and cloned by VectorBuilder (VectorBuilder Inc). All constructs utilized the pmRVac1 backbone and incorporated a T7 promoter-driven IVT cassette, an optimized 5′ untranslated region (UTR) containing a Kozak sequence, a specified open reading frame (ORF), a human α-globin 3′ UTR, and a segmented poly(A) tail (∼110 nt) to ensure efficient transcription and mRNA stability. The EGFP control construct encoded only EGFP, whereas KCNA3-containing constructs encoded full-length human KCNA3 and EGFP, linked by a self-cleaving peptide (P2A) to facilitate multicistronic expression. All plasmids were propagated in E. coli, sequence-verified by Sanger sequencing, and validated by restriction digestion prior to delivery. Glycerol stocks of the verified E. coli clones were provided by VectorBuilder and served as the source material for plasmid preparation and subsequent IVT reactions.

### In vitro transcription and mRNA capping

Plasmid DNA templates encoding KCNA3 and EGFP or EGFP only (Fig. 2C) were linearized using the BspQI restriction enzyme (New England Biolabs) following the manufacturer’s protocol. Briefly, 1 µg of plasmid DNA was digested and purified using the Monarch PCR & DNA Cleanup Kit (New England Biolabs) following the manufacturer’s protocol. The purified linearized DNA was quantified using a NanoDrop spectrophotometer (Thermo Fisher Scientific). All RNA-related procedures were conducted under RNase-free conditions using nuclease-free reagents, consumables, and work surfaces. IVT reactions were performed according to the manufacturer’s protocol with 1 µg of linearized DNA template and the HiScribe T7 High Yield RNA Synthesis Kit (New England Biolabs), which employs T7 RNA polymerase–mediated transcription to produce mRNA. After IVT, the synthesized mRNA was purified using Monarch Spin RNA Cleanup Kit (New England Biolabs) according to the manufacturer’s protocol and quantified using NanoDrop spectrophotometry. RNA integrity and size distribution were evaluated using the Agilent 2100 Bioanalyzer (Agilent Technologies). One-step enzymatic capping of IVT mRNA was then performed using Vaccinia capping enzyme (10 U/µL, New England Biolabs) in conjunction with mRNA Cap 2′-O-Methyltransferase (50 U/µL, New England Biolabs) and RNase inhibitor (New England Biolabs), following the manufacturer’s instructions to produce Cap 1– modified mRNA. The capped mRNA was then purified and quantified using NanoDrop spectrophotometry, and its quality was assessed using an Agilent 2100 Bioanalyzer. High-quality RNA was characterized by an RNA Integrity Number (RIN) ranging from 8 to 10, as determined by the bioanalyzer, indicating minimal degradation (Fig. S5). Additional criteria included the presence of distinct 18S and 28S ribosomal peaks in the bioanalyzer electropherograms (Fig. S5), as well as spectrophotometry results showing an A260/A280 ratio of approximately 2.0 and an A260/A230 ratio exceeding 1.8. Final capped mRNA preparations were aliquoted to prevent repeated freeze–thaw cycles and stored at −80°C until needed for downstream LNP formulation and functional studies. The IVT and capping workflow are summarized in Fig. S4.

### Nanoparticles

The LNPs were synthesized and functionalized according to a previously established protocol (*21–23*). Briefly, lipid vesicles were prepared by combining chloroform solutions of 1,2-dioleoyl-3-trimethylammonium-propane (DOTAP), phosphatidylcholine (PC), phosphatidylethanolamine-polyethylene glycol-biotin (PE-PEG-biotin), and cholesterol (Avanti Polar Lipids Inc.) at a 5:2:1:1 (DOTAP: PC: PE-PEG-biotin: cholesterol) molar ratio. The mixture was dried under nitrogen gas, rehydrated with phosphate-buffered saline (PBS) at pH 7.4, and subsequently subjected to sonication and extrusion with a polycarbonate 100 nm membrane filter (Whatman Nucleopore filter, Cytiva Life Sciences) using a LIPEX™ Thermobarrel Extruder (Northern Lipids Inc.) to produce unilamellar vesicles with a diameter of 100 nm. The vesicle size and polydispersity index (PDI) were verified using a Zetasizer Nano ZS (Malvern Instruments Inc.). The LNPs were then functionalized to specifically target CD5^+^ T cells by conjugating them with a biotinylated anti-human CD5 primary antibody (10 μg/mL) or biotin mouse IgG1 isotype control antibody (10 μg/mL) and Alexa 647-conjugated streptavidin (SAV, Biolegend). More details on these functionalizing antibodies, including clone information and RRIDs, are provided in Table S2. Following purification on Sepharose CL-4B columns (GE Healthcare Bio-Sciences AB) to remove the unbound antibodies, the LNPs were frozen overnight at -80°C and lyophilized overnight (FreeZone2.5 lyophilizer, Labconco) and stored at –80°C. Subsequently, lyophilized LNPs were reconstituted in 0.25 mg/ml of either KCNA3 mRNA (KCNA3-LNPs) or non-targeting EGFP mRNA (EGFP-LNPs) in conjunction with protamine sulfate (MilliporeSigma) at a 1:5 molar ratio in 100 μl of nuclease-free water. The particle concentration of the resulting formulation was quantified by nanoparticle tracking analysis (NanoSight NS300, Malvern Instruments Inc) and measured at 7.19 ± 1.51 x 10⁸ particles/ml.

### mRNA encapsulation efficiency measurement

EGFP mRNA encapsulation efficiency within the LNPs was assessed using the Quant-iT RiboGreen RNA assay kit (ThermoFisher Scientific) following a modified version of the manufacturer’s protocol. This fluorescence-based assay relies on the differential accessibility of the RiboGreen dye to mRNA, which fluoresces brightly only when it binds to accessible RNA in solution. LNP samples were prepared in duplicate: one set was treated directly with the RiboGreen reagent to quantify "free" (non-encapsulated) mRNA, while the other set was first treated with 0.5% Triton X-100 detergent (MilliporeSigma) to disrupt the LNPs and release all encapsulated mRNA for quantification of "total" mRNA. Fluorescence measurements were conducted using a Biotek Synergy H1 plate reader (excitation 485 nm, emission 528 nm), and concentrations were interpolated from a standard curve generated using known concentrations of unencapsulated mRNA.

### Human samples and subjects

For this study, discarded blood units from the Hoxworth Blood Center at the University of Cincinnati were utilized. Age and demographic data of these units were unavailable. The study protocols and informed consent forms were approved by the University of Cincinnati Institutional Review Board (IRB no. 2013-2516).

### Cell isolation and in vitro activation

PBMCs were isolated from whole blood using Ficoll Paque density gradient centrifugation (GE Healthcare Biosciences), as previously described (*23*). CD3 ^⁺^ T cells were subsequently isolated from PBMCs via negative selection using the EasySep Human T Cell Enrichment Kit (STEMCELL Technologies Inc.), whereas CD8 ^⁺^ T cells were isolated via negative selection from PBMCs using the EasySep Human CD8 ^⁺^ T Cell Enrichment Kit (STEMCELL Technologies Inc.) according to the manufacturer’s protocol. Cells were cultured in RPMI 1640 medium (ThermoFisher Scientific) supplemented with 10% human serum (Millipore Sigma), penicillin (200 U/mL), streptomycin (200 μg/mL), 1 mM l-glutamine, and 10 mM HEPES (all from ThermoFisher Scientific) (*22*).

### T cell activation

T cells were activated using plate-bound mouse anti-human CD3 and CD28 antibodies, each at a concentration of 10 μg/mL (Biolegend), as previously described (*22, 23*). The activation times varied by experiment and are presented in the individual figure legends.

### In vitro LNP treatment

For in vitro experiments, 2 x 10⁵ PBMCs or T cells were mixed with 50 μl of KCNA3- or EGFP-LNPs (7.19 x 10^8^ particles/ml; input mRNA concentration, 0.25 mg/ml) in 250 μl of T cell medium, resulting in a final volume of 300 μl. This resulted in 3.60 x 10^7^ particles per well, a final particle concentration of 1.20 x 10^8^ particles/ml, a 1:180 cell-to-particle ratio, and approximately 12.5 μg of input mRNA per well, yielding a final encapsulated mRNA concentration of 41.67 μg/ml.

### Flow cytometry

The cell specificity of LNPs functionalized with the anti-CD5 antibodies (CD5-LNPs) was determined by measuring their association with immune cell subsets using flow cytometry. Surface staining and immunophenotyping for flow cytometry were conducted according to established protocols. Briefly, HD PBMCs were treated with either Alexa Fluor 647-conjugated CD5-LNPs or IgG-LNPs (LNPs functionalized with IgG isotype control antibody) for 24 h. 1 × 10^6^ cells per condition were then stained with Zombie Red Live/Dead stain and the following antibodies in Cell Staining buffer (Biolegend) according to the manufacturer’s protocol: Brilliant Violet (BV) 421-conjugated mouse anti-human CD3, BV 785-conjugated mouse anti-human CD4, BV 711-conjugated mouse anti-human CD8, APC-CY7-conjugated mouse anti-human CD19, PE-Dazzle 594-conjugated mouse anti-human CD56 and PE-conjugated mouse anti-human CD14. The antibody details, including clone information and RRIDs, are provided in Table S2. After staining, cells were washed, fixed in 4% paraformaldehyde, and resuspended in staining buffer. Data were acquired using a Cytek Northern Lights Spectral Flow Cytometer (Cytek Biosciences). Daily instrument quality control and performance tracking were performed using Cytek Assay Settings beads according to the manufacturer’s instructions. Spectral unmixing was performed using single-stained reference controls and the SpectroFlo software (Cytek Biosciences). Forward and side scatter parameters were used to exclude debris, and doublets were excluded using FSC-A versus FSC-H gating. Viable cells were identified using the Zombie Red Live/Dead Stain. The LNP population (Alexa Fluor 647^+^ population) was identified within the various immune cell subsets for both the CD5- and IgG-LNPs. Data were analyzed using FlowJo software (BD Biosciences), and the gating strategies are illustrated in Fig. S6. The gating strategy has been described previously by us (*27*).

To assess mRNA delivery into cells, HD PBMCs were activated and simultaneously treated for 48 h with Alexa Fluor 647-conjugated EGFP-LNPs or KCNA3-LNPs. Cells treated with LNPs without mRNA (empty-LNPs) and those not treated with LNPs (no LNPs) served as controls. 1 × 10^6^ cells per condition were then washed with PBS and stained with Zombie UV Live/Dead stain. Cells were washed with cell staining buffer (Biolegend) and fixed with 4% paraformaldehyde for 30 min. Subsequently, cells were stained with surface guinea pig anti-Kv1.3 (Alomone Labs) primary antibody overnight at 4°C. The following day, cells were washed twice in cell staining buffer, followed by incubation with a secondary anti-guinea pig (Alexa Fluor 555 goat anti-guinea pig IgG; ThermoFisher) antibody. The cells were then stained with an antibody cocktail consisting of BV 605 conjugated mouse anti-human CD3, and BV 711 conjugated mouse anti-human CD69 antibodies. The complete details of the antibodies used in these sets of experiments are presented in Table S2. We have previously confirmed the specificity of the Kv1.3 antibody (*14*). Data were acquired using a BD Fortessa flow cytometer (BD Biosciences) and analyzed using FlowJo software (BD Biosciences). Compensation was performed using unstained and single-stained PBMCs from HDs, along with UltraComp eBeads (ThermoFisher). EGFP expression and the abundance of Kv1.3 and CD69 were measured within the CD3^+^LNP^+^ populations. The gating strategy used in these experiments is illustrated in Fig. S7.

### Microscopy

To assess mRNA delivery into the cells, HD PBMCs were activated and simultaneously treated for 48 h with Alexa Fluor 647-conjugated EGFP-LNPs or KCNA3-LNPs. Cells treated with LNPs lacking mRNA (empty LNPs) and those not treated with LNPs (no LNPs) were used as controls. After 48 h, the cells were visualized for LNP and EGFP fluorescence using a Zeiss 710 laser scanning confocal microscope (Zeiss GmBH) with a 20× lens at room temperature, employing a pinhole setting of 1 airy unit. Data were collected using the microscope’s “Multi-Track” option to minimize channel crosstalk, and more than five fields per well were imaged.

### Electrophysiology

K^+^ currents were recorded using a whole-cell patch clamp configuration with an AxoPatch 200B amplifier (Molecular Devices). The external solution contained (in mM): 150 NaCl, 5 KCl, 2.5 CaCl2, 1 MgCl2, and 10 HEPES, at pH 7.4. The pipette solution consisted of (in mM): 140 KF, 1 CaCl2, 10 EGTA, 2 MgCl2, and 10 HEPES, pH 7.4 (*21*). The K^+^ current was measured in voltage-clamp mode and induced by step-pulse depolarization to +40 mV from −80 mV holding potential (HP), 800 ms duration, every 30 s. The digitized signals were stored and analyzed using the pClamp 9 software (Molecular Devices).

### Intracellular Ca^2+^ measurement

Intracellular Ca^2+^ flux was assessed using the Ca^2+^ add-back method, as previously described (*14, 21*). Briefly, freshly isolated CD8⁺ T cells were treated with KCNA3-LNPs or EGFP-LNPs (1.20 x 10^8^ particles/ml; 41.67 μg/ml encapsulated mRNA) and activated with anti-CD3/CD28 antibodies for 48 hours. Cells (1 x 10⁶) were loaded with the ratiometric Ca^2+^ indicator Indo-1 AM (2 mg/ml; 1:1000 dilution; ThermoFisher) in the presence of 0.015% Pluronic F-127 (ThermoFisher). The cells were incubated in Hank’s Balanced Salt Solution (HBSS) supplemented with 1 mM CaCl₂, 1 mM MgCl₂, and 1% FBS (ThermoFisher) for 30 min at 37°C. Following dye loading, the cells were washed three times with HBSS containing 10 mM HEPES (pH 7.0; ThermoFisher) and 1% FBS. Prior to acquisition, the cells were resuspended in Ca^2+^-free HBSS/HEPES buffer containing 0.5 mM EGTA (pH 7.4) and maintained at 37°C. Real-time Indo-1 fluorescence ratios (violet/blue emission) were acquired using a BD LSRFortessa flow cytometer with previously optimized instrument settings. To assess Ca^2+^ influx, cells were first stimulated with thapsigargin (1 μM, MilliporeSigma) in Ca^2+^-free buffer to deplete intracellular Ca^2+^ stores and activate CRAC channels. This was followed by the addition of 2 mM extracellular Ca^2+^, allowing Ca^2+^ to enter the cells through open CRAC channels. Data were analyzed using FlowJo software (BD Biosciences), and intracellular Ca²⁺ flux was quantified as the fold change in the Indo-1 fluorescence ratio, calculated by dividing the post– Ca^2+^ addition peak by the baseline fluorescence in Ca²⁺-free conditions.

### IFN-γ measurements

HD CD8^+^ T cells were treated with KCNA3- and EGFP-LNPs at final concentrations of 0.6 x 10^8^, 1.20 x 10^8^ and 2.4 x 10^8^ particles/ml, and 20.83 μg/ml, 41.67 μg/ml and 83.34 μg/ml encapsulated mRNA. The cells were activated with anti-CD3/CD28 antibodies for 48 h. In a separate set of experiments, HD CD8^+^ T cells were treated with KCNA3-EGFP- and empty-LNPs at a final LNP concentration of 1.20 x 10^8^ particles/ml and encapsulated mRNA concentration of 41.67 μg/ml (CD8^+^ T cells without LNPs served as controls) and activated with anti-CD3/CD28 antibodies for 48 h. IFN-γ levels in cell culture supernatants were measured using a commercially available ELISA kit (ThermoFisher), following the manufacturer’s instructions.

### LEGENDplex™ Cytokine and Cytotoxicity Assay

Cytokine and cytotoxic marker secretion by CD8^+^ T cells was assessed using the LEGENDplex Human CD8/NK Panel (BioLegend; Cat# 741187), which quantified IL-17A, IL-2, IL-6, TNF-α, soluble Fas (sFas), soluble Fas ligand (sFasL), IFN-γ, Granzyme A, Granzyme B, Perforin, and Granulysin. The assay was performed according to the manufacturer’s instructions. Briefly, HD CD8^+^ T cells were activated with anti-CD3/CD28 antibodies for 48 h in the presence of KCNA3- LNPs, EGFP-LNPs (1.20 x 10^8^ particles/ml; 41.67 μg/ml encapsulated mRNA), empty LNPs (1.20 x 10^8^ particles/ml) or no LNPs. Supernatants (25 μl) were collected and the assay was performed in duplicate in a 96-well plate according to the manufacturer’s instructions. Samples were acquired on a Cytek Aurora spectral flow cytometer (Cytek Biosciences) equipped with a yellow-green laser and an automated 96-well plate reader using SpectroFlo software (Cytek Biosciences) and manufacturer-recommended settings. Data were analyzed using the LEGENDplex™ Data Analysis Software (Biolegend). The results are reported as a fold change in the mean fluorescence intensity (MFI) normalized to the no LNP control group.

### Cal 27 cell culture

Cal 27 cells, obtained from the American Tissue Culture Collection (ATCC, Manassas, VA, USA), were cultured in Dulbecco’s modified Eagle’s medium (DMEM) supplemented with 10% fetal bovine serum, 1% penicillin–streptomycin, 1% minimum essential amino acids, 6 mM L-glutamine, and 1% sodium pyruvate, as previously described (all reagents from ThermoFisher) (*52*). Cell cultures were maintained at 37°C in a humidified incubator with 5% CO2. Mycoplasma testing was routinely performed using the My-coStrip Mycoplasma Detection Kit (Invivogen, Catalog# rep-mys-50) and yielded negative results for all cultures.

### Humanized PBMC-HNSCC xenograft mouse model generation and validation

NSG mice were obtained from the Comprehensive Mouse and Cancer Core (Division of Experimental Hematology and Cancer Biology) at Cincinnati Children’s Hospital Medical Center (CCHMC). Six-week-old mice of both sexes were used in the experiments, and their flanks were injected with a Cal27 spheroid suspension. Cal27 human oral squamous carcinoma cells were cultured as 3D spheroids using a Matrigel overlay method. Cells were seeded onto Matrigel-coated wells to promote 3D growth, as previously described (*53*). Briefly, 20,000 cells were seeded into 24-well plates coated with Matrigel (Corning) and maintained in a medium containing 1% Matrigel for four days. Spheroids were harvested by adding 1 ml of ice-cold Cell Recovery Solution (Corning) per well, incubating at room temperature for 5 min, gently pipetting to disrupt the Matrigel, and transferring the plates to 4°C for 20 min. Matrix dissolution and spheroid release were confirmed microscopically. All materials were collected into 50 ml tubes, the wells were rinsed with PBS, and the remaining spheroids were recovered by repeating the dissociation steps as needed. Spheroids were pelleted at 300 × g for 5 min and pooled.

To determine the number of cells delivered per mouse, an aliquot of the pooled spheroids was dissociated with TrypLE (ThermoFisher) for 15 min at 37°C. Complete dissociation was confirmed using microscopy. Cell counts were consistent across measurements (2.12–2.46 × 10⁵ cells/ml; 96–98% viability), and an average concentration of 2 × 10⁵ cells/mL was used for dosing calculations. Each mouse received 200 μl of a 1:1 PBS:Matrigel spheroid suspension, corresponding to ∼4 × 10⁴ Cal27 cells per injection. When the tumors were palpable (23 days post-spheroid injection), each mouse received 10 × 10^6^ human PBMCs in 0.5 ml PBS s.c. on the perimeter of the tumor but not inside. The mice were assigned to four groups: Cal27 + PBMCs + KCNA3-LNPs (n = 7), Cal27 + PBMCs + EGFP-LNPs (n = 7), Cal27 + PBMCs + empty LNPs (n = 5), and Cal27 + PBMCs only (n = 2). Mice received 100 μl of their designated LNPs formulation (7.19 x 10^8^ particles/ml) per injection, at each time point by subcutaneous injection within 5 mm proximity to the tumor around the tumor but not inside the tumor. HD PBMCs from a single donor were administered on day 24 post-spheroid injection and LNPs were administered on days 24, 28, 31, 35, 38, and 40 (Fig. 7A). On Day 41, all animals were euthanized via CO2 inhalation and sacrificed for tumor excision and tissue (tumor, spleen, liver, lung and kidneys) collection. The length (L), width (W) and height (H) of the tumor were measured using calipers, and tumor volume (V) was calculated using the formula *V* = <u>^π^</u> *x* (*L x W x H*). The study protocol was approved by the Institutional Animal Care and Use Committee (IACUC) of CCHMC (IACUC protocol no. 2021-0032), and the University of Cincinnati (protocol no. 21-10-29-01), and all efforts were made to minimize animal suffering. All mouse experiments were performed in accordance with the relevant guidelines and regulations. This study adhered to the Animal Research: Reporting of In Vivo Experiments (ARRIVE) guidelines 2.0, specifically regarding detailed descriptions of animal characteristics, experimental procedures, and statistical methods.

### Immunohistochemistry and histopathologic evaluation

Excised HNSCC mouse tumors were formalin-fixed, paraffin-embedded (FFPE), and sectioned at a thickness of 5 μm at the Integrated Pathology Research Facility (IPRF) at CCHMC. The sections were stained using standard IHC protocols. Briefly, slides were deparaffinized and stained with monoclonal rabbit anti-human CD8 (clone SP57, Roche) and monoclonal mouse anti-human Granzyme B (cat# MA1-35461, ThermoFisher Scientific) using a Ventana BenchMark ULTRA automated IHC slide staining system (Roche), as previously described (*23, 54*). For dual CD8 and Granzyme B staining, primary antibodies were indirectly detected using the ultraView Universal Alkaline Phosphatase Red detection kit (Roche) and the ultraView Universal DAB Detection Kit (Roche), which includes a horseradish peroxidase (HRP) multimer and 3,3′-diaminobenzidine tetrahydrochloride (DAB) chromogen. The slides were counterstained with hematoxylin. IHC images were acquired at 20× magnification using a Leica DMi8 microscope equipped with the Application Suite X software (Leica Microsystems Inc.). Five representative fields were selected from each slide for imaging. Image analysis was performed using the ImageJ software, as previously described (*23, 54*). Within each field, a region of interest (ROI) was delineated around the tumor area using the manual drawing tool in ImageJ. The number of CD8^+^ T cells (indicated by the brown signal) within each ROI was quantified, normalized to the ROI area, and expressed as cells per square millimeter. For dual staining, we quantified the CD8^+^ T cell signal (brown) with the Granzyme B signal (red) in each ROI using ImageJ, expressing them as a percentage of Granzyme B^+^CD8^+^ T cells.

For histological evaluation, additional FFPE sections were stained with H&E at the CCHMC IPRF using standard protocols. H&E-stained slides were initially reviewed by a board-certified pathologist to assess overall histopathology across all experimental groups via light microscopy at 40× magnification. Digital images were acquired using an Olympus DP23 CMOS camera mounted on an Olympus microscope system. Entire tumor sections were evaluated, and intratumoral lymphocyte infiltration (TIL) was subsequently scored in a separate blinded analysis, with group allocation concealed from the pathologist. TIL infiltration was scored semi-quantitatively on a scale of 0–3, where 0 represented minimal or no infiltration, 1+ represented mild infiltration, 2+ represented intermediate infiltration, and 3+ represented severe infiltration. The histopathological scores are summarized in Table S1.

## Statistical analysis

Statistical analyses were conducted between groups using the Student’s t-test (paired or unpaired) when appropriate. When equality of variances could not be assumed, Welch’s correction was applied to the unpaired t-test. For non-normally distributed data, comparisons between two groups were performed using the Mann–Whitney U test. For comparisons between multiple groups, one-way analysis of variance (ANOVA) was performed if the data were normally distributed, or the Kruskal–Wallis test was used if normality was not assumed. Post-hoc analyses following ANOVA or non-parametric tests utilized Holm-Sidak, Tukey, or Dunn’s methods for multiple pairwise comparisons, as appropriate. Normality of sample distributions was assessed using the Shapiro– Wilk test. All statistical tests were performed using GraphPad Prism 10.6.1 (GraphPad Software LLC, Boston, MA, USA). Statistical significance was set at P ≤ 0.05 was considered statistically significant. Detailed information regarding specific statistical tests and their corresponding significance values can be found in the individual figures and legends.

## List of Supplementary Materials

Fig.S1. Association of KCNA3 expression with overall survival in TCGA clear cell renal cell carcinoma (ccRCC).

Fig. S2. Synthesis and antibody labeling of unilamellar lipid vesicles (ULVs).

Fig. S3. Particle size distribution of ULVs.

Fig. S4. In vitro transcription (IVT) workflow for mRNA production.

Fig. S5. RNA quality assessment of in vitro-transcribed KCNA3 mRNA.

Fig. S6. Flow cytometric gating strategy for identifying major peripheral blood immune cell populations.

Fig. S7. Flow cytometry analysis of KCNA3-LNP uptake and expression of Kv1.3 and CD69 in human PBMCs.

Fig. S8. IFN-γ levels in CD8+ T cells treated with LNPs.

Fig. S9. Comparison of tumor volumes in EGFP-LNP- and no LNP-treated hu-PBMC-HNSCC mice.

Fig. S10. Weights of the livers, lungs, and kidneys dissected from hu-PBMC-HNSCC mice injected with EGFP-LNPs or KCNA3-LNPs.

Fig. S11. Histopathological evidence of enhanced inflammatory infiltration and tumor cell death in hu-PBMC-HNSCC tumors treated with KCNA3-LNPs.

Fig. S12. Comparison of immune cell infiltration in EGFP-LNP-injected hu-PBMC-HNSCC mice and mice that did not receive LNPs.

Table S1 Semiquantitative histopathology analysis of intratumoral lymphocyte infiltration in hu-PBMC-HNSCC mice

Table S2 Antibodies used for LNP functionalization and Flow Cytometry experiments

## Acknowledgments

The authors are grateful to Dr. Roman Jandarov (Department of Biostatistics and Bioinformatics, University of Cincinnati) for his valuable advice on statistical analysis and for reviewing all statistical analyses. We also thank the Department of Pharmacology, Physiology and Neurobiology, University of Cincinnati, for providing the Zetasizer Nano ZS. Confocal microscopy images were acquired at the Live Microscopy Core (Department of Pharmacology and Systems Physiology, University of Cincinnati). We also thank Dr. Takahisa Nakamura (Division of Endocrinology, CCHMC) for generously providing access to the NanoSight NS300 instrument. All flow cytometry experiments were performed using equipment maintained by the Research Flow Cytometry Facility (Division of Rheumatology) at CCHMC. RNA quality control was performed at the Genomics, Epigenomics and Sequencing Core, Department of Environmental Health, University of Cincinnati. Immunohistochemistry and histological staining were performed at the Integrated Pathology Research Facility, CCHMC. The experimental schema were drawn using BioRender.

## Funding sources

This research was supported by a grant from the National Institutes of Health (R21CA277341) and Dean Bridge funds, as well as a Research Innovation/Pilot Grant from the University of Cincinnati College of Medicine, and a Marlene Harris Ride Cincinnati Breast Cancer Research Fund pilot grant from the University of Cincinnati Cancer Center, all awarded to Dr. Laura Conforti. Flow cytometry experiments conducted at the Research Flow Cytometry Facility, CCHMC were supported by an NIH grant (AR070549). Stipend support for A.I. was provided by the UC MedEx SURF program and divisional funds from the Division of Nephrology at the University of Cincinnati.

## Author contributions

Conceptualization: AAC, LC Methodology: AAC, MVK, LC

Investigation: AAC, MVK, AI, AOA, SS, HSN, BHH Visualization: AAC

Funding acquisition: LC

Project administration: AAC, MVK, LC Supervision: LC

Writing – original draft: AAC, LC

All authors discussed the results and commented on the manuscript.

## Competing interests

The authors declare no competing interests

## Data and materials availability

All data associated with this study are available in the main text or supplementary materials.

## Notes

### Competing Interest Statement

The authors have declared no competing interest.

