## Supplemental Material for "Ion channel-enhancing nanoparticles reprogram T cells in vivo for cancer immunotherapy"

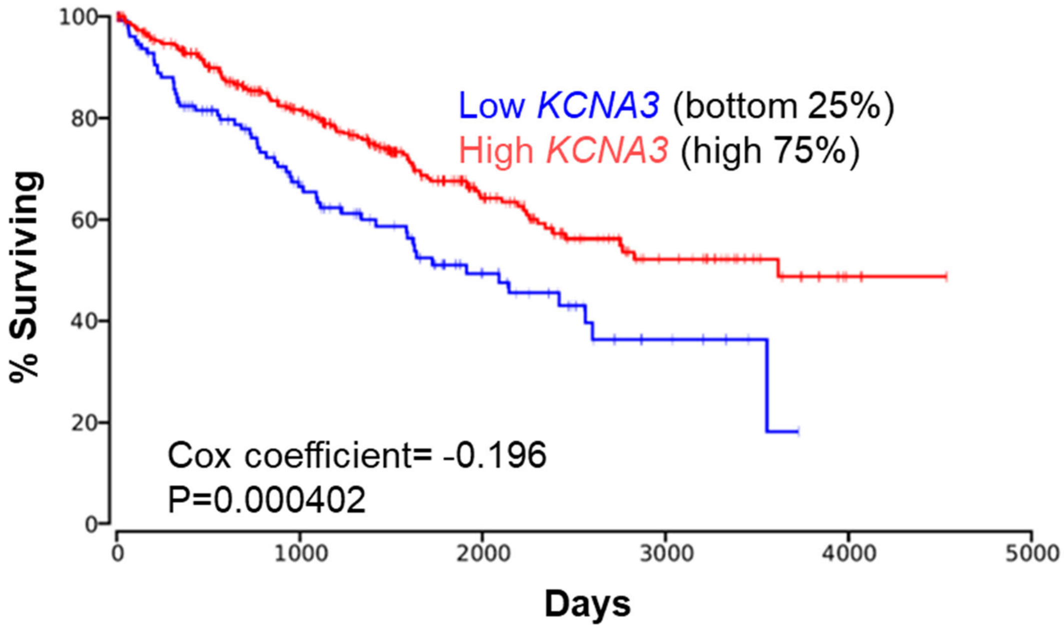


**Fig. S1. Association of KCNA3 expression with overall survival in TCGA clear cell renal cell carcinoma (ccRCC).** Kaplan-Meier analysis of overall survival in the TCGA ccRCC cohort, stratified by KCNA3 expression. Patients were divided into KCNA3-low (bottom 25%) and KCNA3-high (top 75%) groups. Higher KCNA3 expression was significantly associated with improved overall survival compared to lower expression. Cox proportional hazards analysis revealed a negative coefficient (–0.196), indicating a protective effect of KCNA3 expression on survival (P = 0.000402).

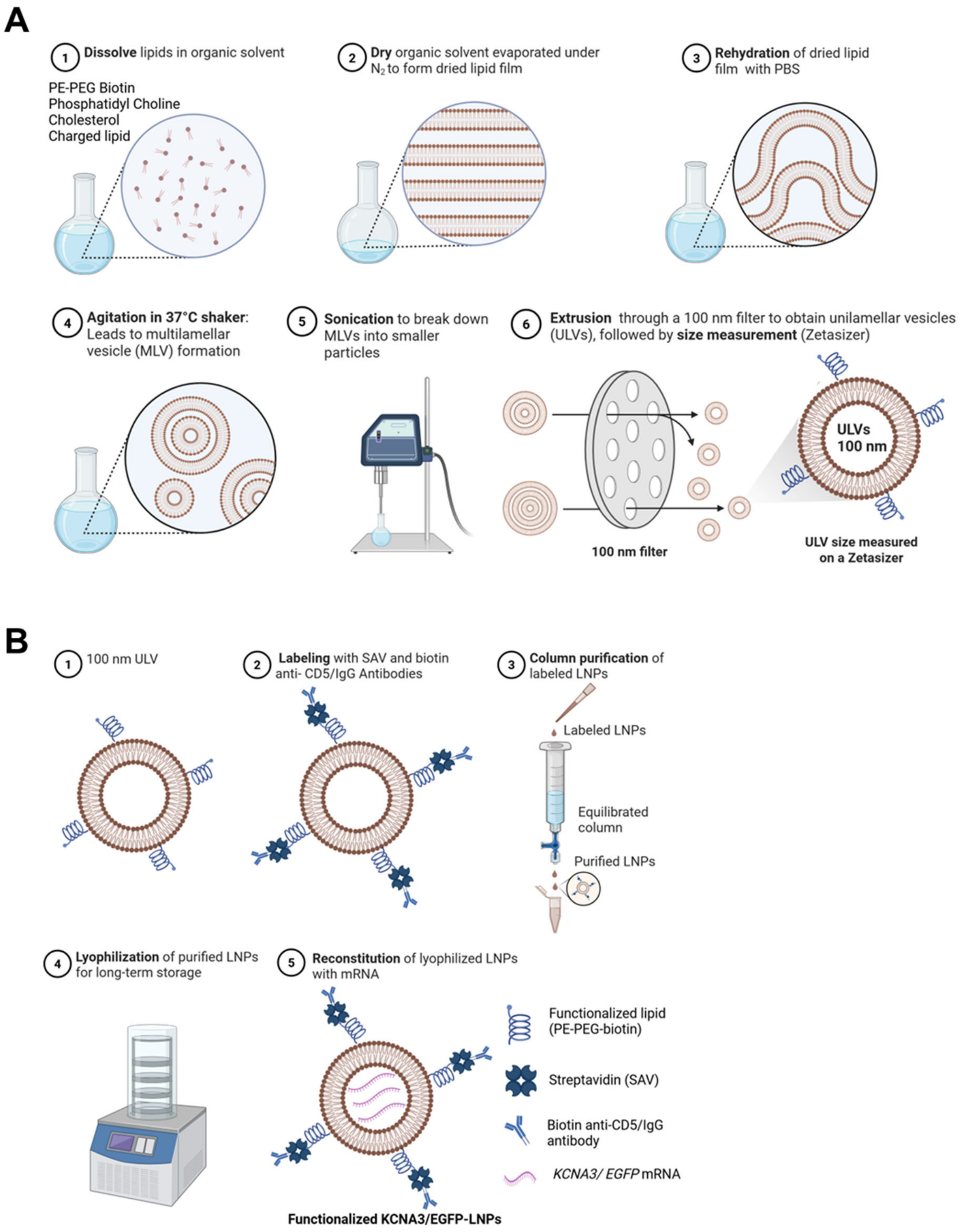


**Fig. S2. Synthesis and antibody labeling of unilamellar lipid vesicles (ULVs).** (A) A schematic of ULV synthesis is shown: lipids were combined in round-bottom glass vessels at the ratios specified in the Materials and Methods, and the resulting film was dried under nitrogen. The lipids were then rehydrated in phosphate-buffered saline (PBS; pH 7.4) and shaken at 37°C for 2 h, followed by sonication and extrusion through 100 nm filter membranes to produce ULVs with an approximate diameter of 100 nm. ULV particle size was determined using a Malvern Zetasizer Nano ZS. (B) ULV functionalization was achieved by attaching biotinylated anti-CD5 antibodies to PE-PEG-biotin via streptavidin binding. The lyophilized antibody-coated nanoparticles were reconstituted with mRNA prior to downstream applications. Detailed protocols are provided in Materials and Methods.

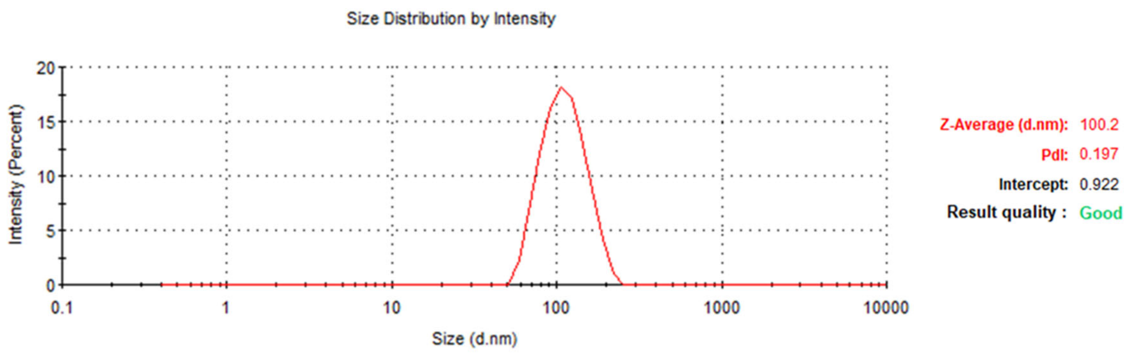


**Fig. S3. Particle size distribution of ULVs.** Representative size distributions of ULVs eluted in sterile-filtered PBS were measured using a Malvern Zetasizer Nano ZS. The results indicate a monodisperse peak with a Z-average diameter of 100.2 nm and a polydispersity index (PDI) of 0.197, signifying a narrow size distribution.

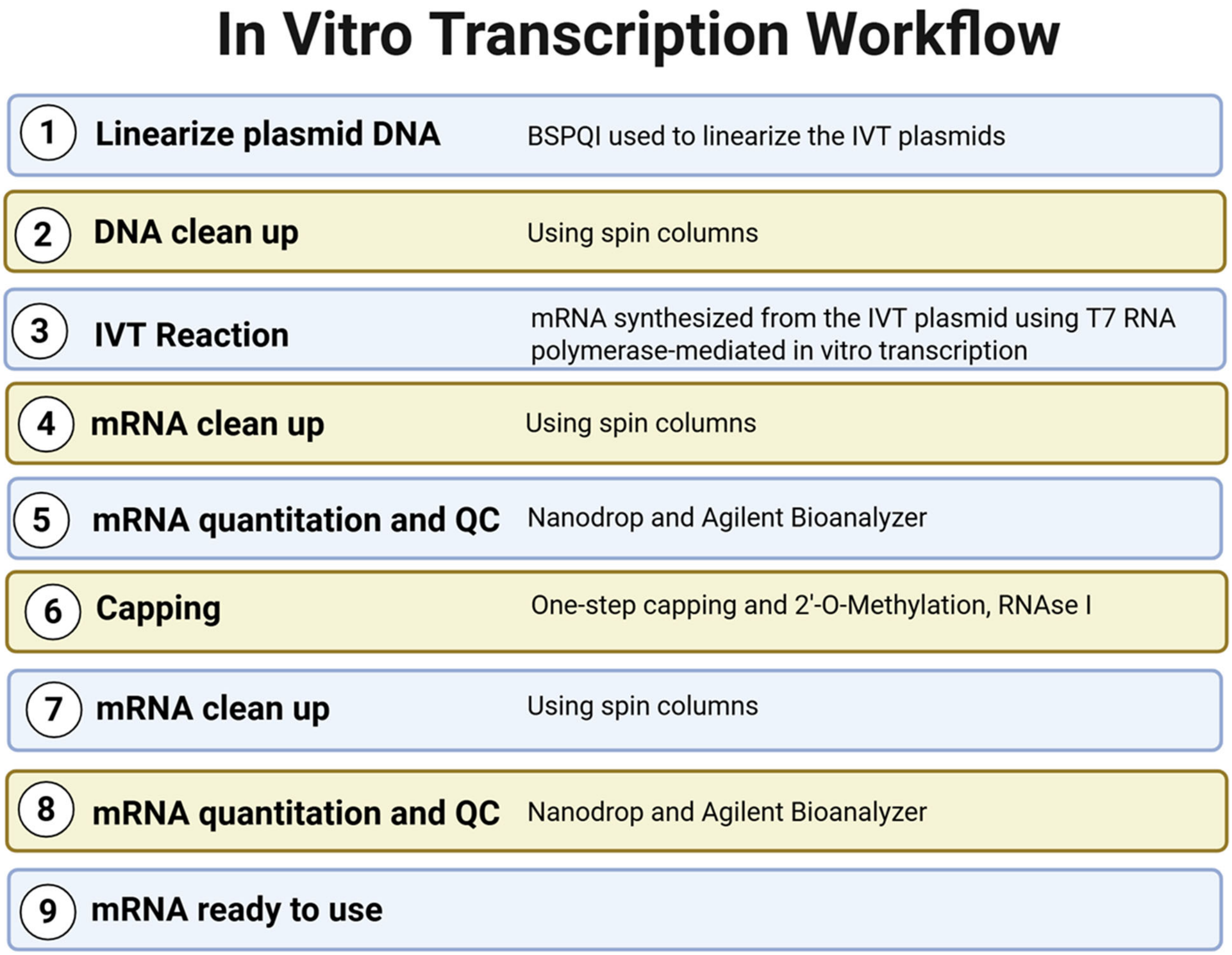


**Figure S4. In vitro transcription (IVT) workflow for mRNA production.** A schematic overview detailing the step-by-step IVT process used to generate capped, high-quality mRNA. Initially, IVT plasmids were linearized with BspQI, followed by DNA cleanup using spin columns. Subsequent in vitro transcription with T7 RNA polymerase synthesized mRNA, which was then purified and evaluated for yield and integrity via NanoDrop spectrophotometry and Agilent Bioanalyzer analysis. Following enzymatic capping and 2′-O-methylation, including RNase I treatment, additional cleanup and quality control steps were performed. Finally, the product was purified using a Monarch Spin RNA

Cleanup Kit and quantified using NanoDrop spectrophotometry. Agilent Bioanalyzer analysis and ratios from spectrophotometry confirmed the suitability of the mRNA for downstream applications (see Materials and Methods for details, and quality control criteria).

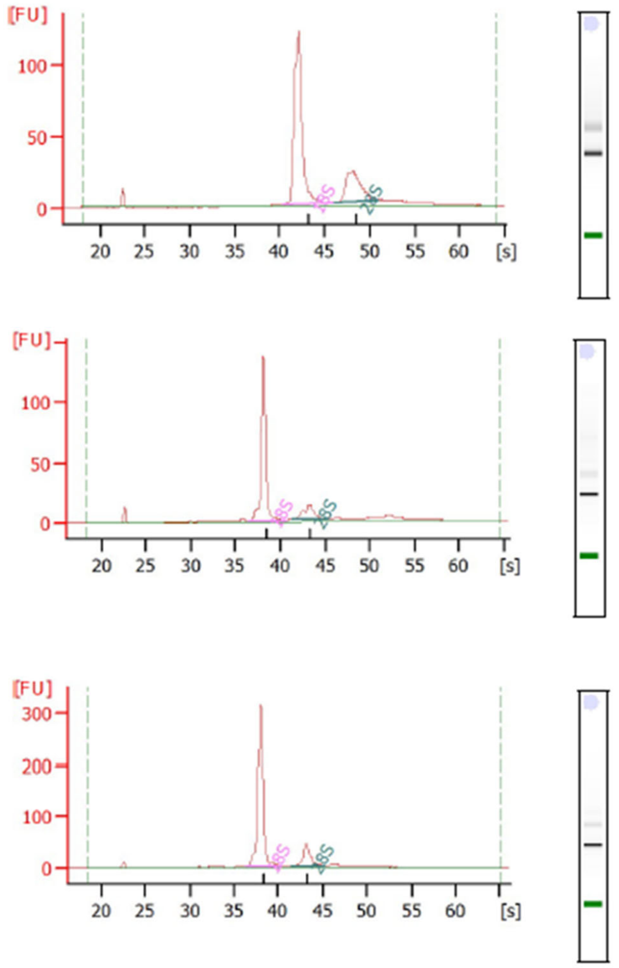


**Fig. S5. RNA quality assessment of in vitro-transcribed KCNA3 mRNA.** Agilent 2100 Bioanalyzer (Eukaryote Total RNA Nano chips) electropherograms (left) and corresponding virtual gel images (right) were generated for three representative KCNA3 mRNA preparations produced via in vitro transcription. Electropherograms display RNA migration time in seconds (s) on the horizontal x-axis and fluorescent signal intensity in fluorescence units (FU) on the vertical y-axis. The 18S and 28S ribosomal peaks are highlighted. All preparations exhibited intact RNA profiles, with RNA Integrity Numbers (RINs ≈ 9.8–10), and demonstrated no evidence of degradation, confirming their suitability for downstream use. (See Materials and Methods for detailed quality control criteria.).

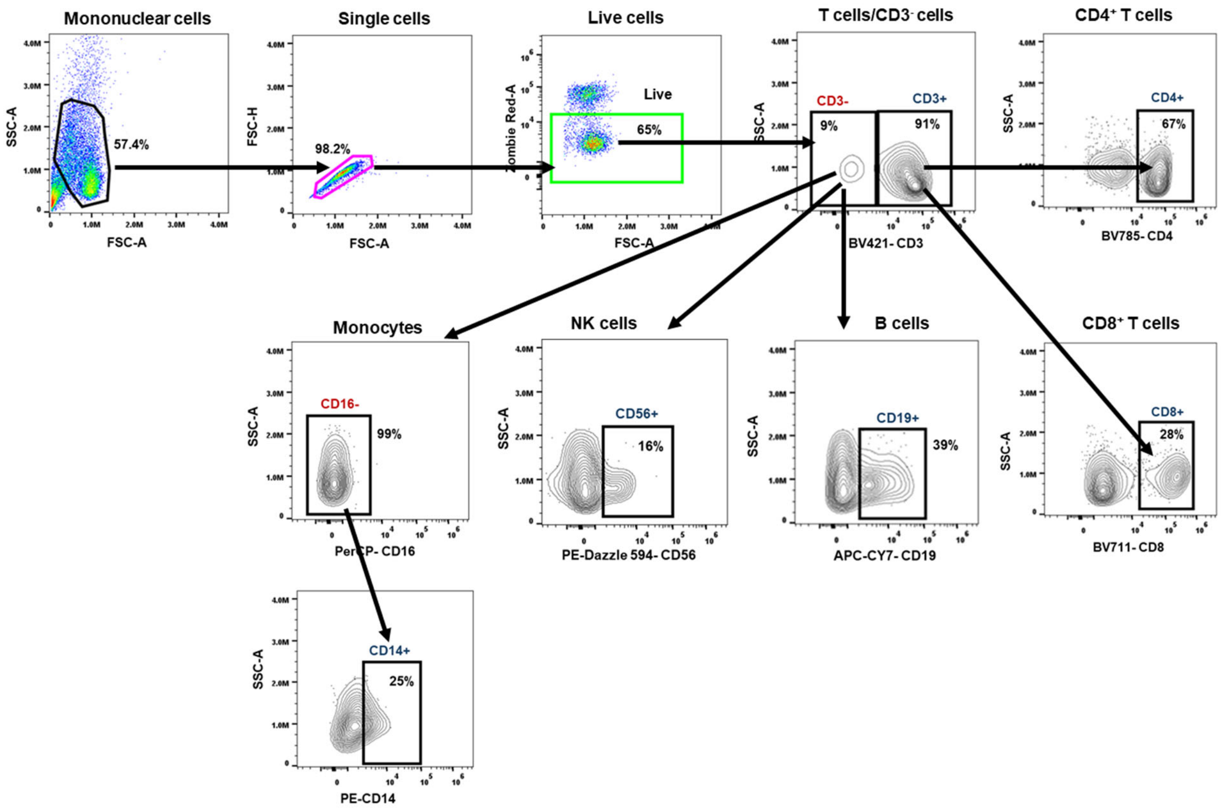


**Figure S6. Flow cytometric gating strategy for identifying major peripheral blood immune cell populations.** Representative plots illustrate a sequential gating approach in which leukocytes were initially selected based on forward and side scatter, followed by the exclusion of doublets and gating on live cells. Live cells were then divided into $CD3^+$ T cells and $CD3^-$ non-T cells. Within the $CD3^+$ population, $CD4^+$ and $CD8^+$ T cell subsets were distinguished. The $CD3^-$ compartment was further subdivided to identify $CD19^+$ B cells, $CD56^+$ NK cells, and $CD16^-$ $CD14^+$ monocytes. The percentages indicated adjacent to each gate represent the proportion of cells within the parent gate. The data are representative of a typical analysis.

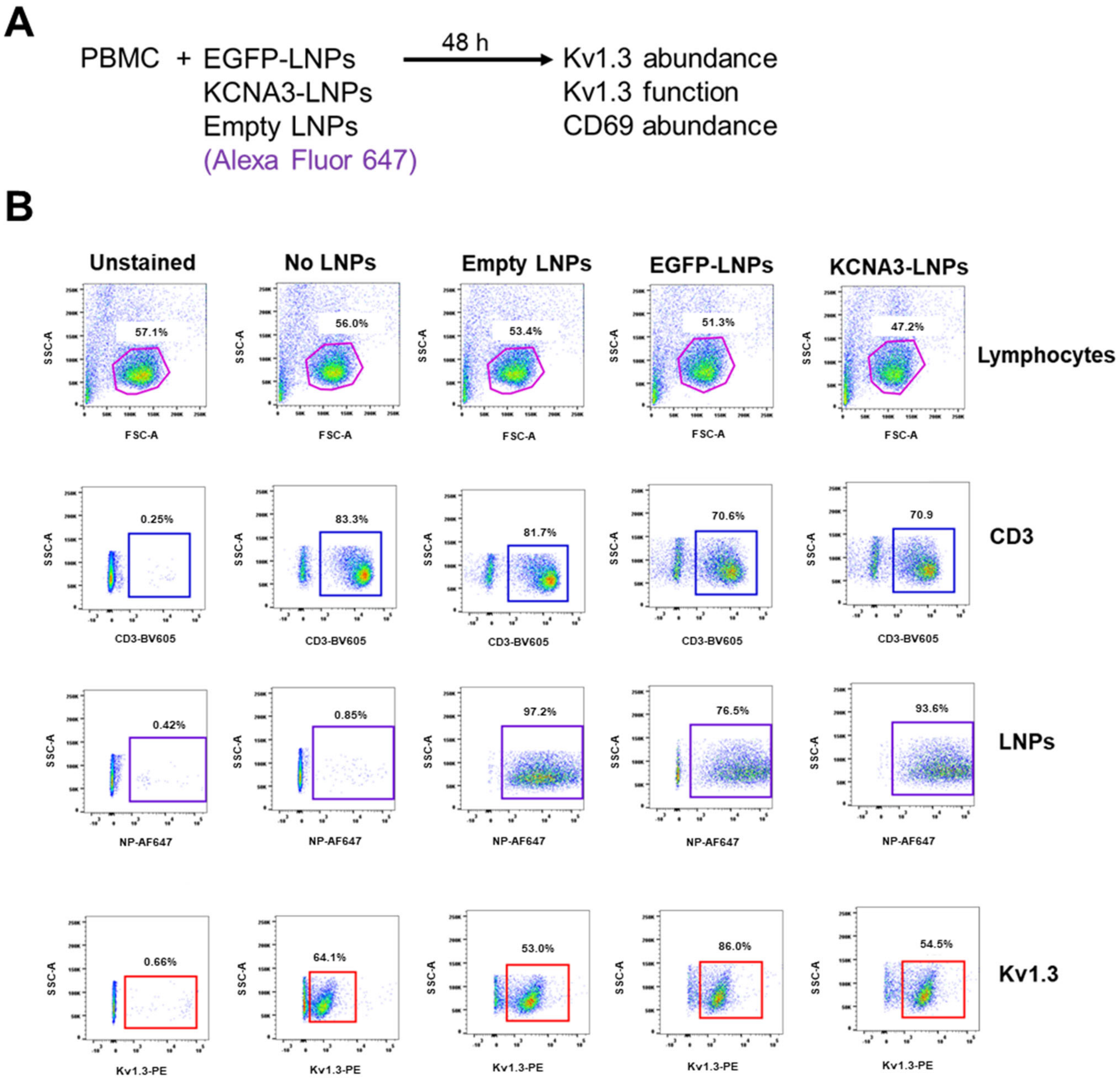


**Figure S7. Flow cytometry analysis of KCNA3-LNP uptake and expression of Kv1.3 and CD69 in human PBMCs.** (A) Experimental protocol for assessing intracellular delivery of KCNA3-LNPs into HD PBMCs and subsequent evaluation of extracellular Kv1.3 and CD69 expression (data shown in Fig. 5). (B) Representative flow cytometry plots illustrate the gating strategy and fluorescence signals used to measure cellular uptake and protein expression following treatment with mRNA-loaded Alexa Fluor 647-labeled LNPs. Lymphocytes were first identified based on forward and side scatter properties. The $CD3^+$ T-cell population was gated within the live cell population, and LNP uptake was assessed within this population. Extracellular Kv1.3 expression (PE signal) was measured in $CD3^+$ cells. The percentage of Kv1.3-positive cells was quantified across the unstained controls, KCNA3-LNPs, EGFP-LNPs, empty LNPs, and no LNP treatment groups. The percentages of Kv1.3-positive cells are indicated above each gate. A similar gating strategy was employed to assess CD69 abundance within the $CD3^+$ population. Finally, gMFI was calculated for Kv1.3 and CD69 populations for all experimental groups (Fig. 4A and B). The data are representative of a typical experiment.

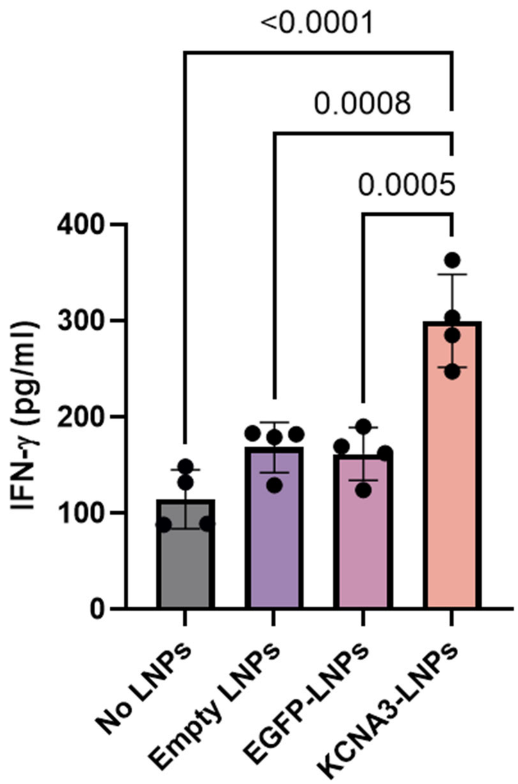


**Figure S8. IFN-γ levels in CD8$^+$ T cells treated with LNPs.** CD8$^+$ T cells from HDs were activated for 48 h and treated with KCNA3-LNPs, EGFP-LNPs (1.20 x $10^8$ particles/ml; 41.67 µg/ml encapsulated mRNA), empty LNPs (1.20 x $10^8$ particles/ml), or left untreated (no LNPs). IFN-γ concentrations were measured in the medium using ELISA. The bars represent the mean ± SD from four HDs. Statistical significance was assessed using one-way ANOVA ($P < 0.0001$), followed by Tukey's post-hoc test. No statistically significant differences were observed between the no LNPs, empty LNPs, and EGFP-LNPs groups.

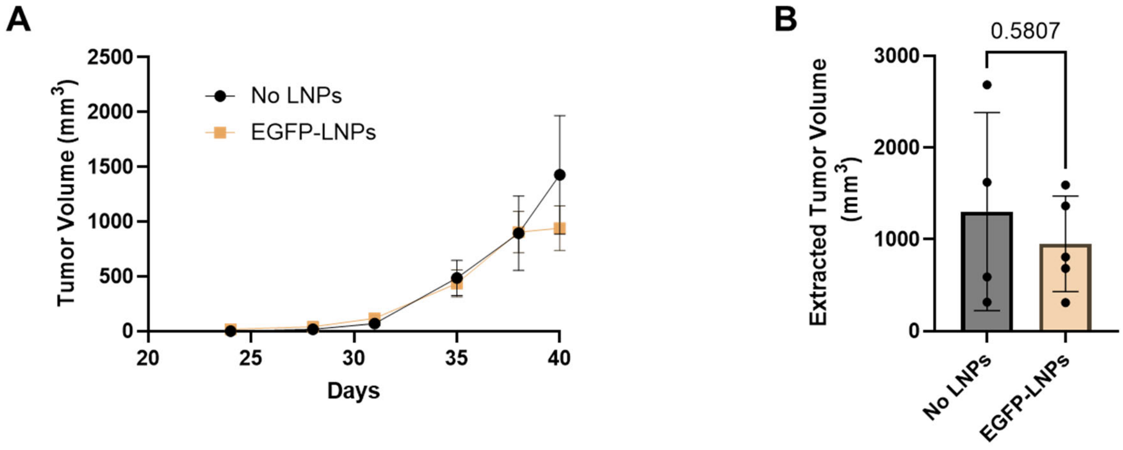


**Figure S9. Comparison of tumor volumes in EGFP-LNP- and no LNP- treated hu-PBMC-HNSCC mice**. (A) Tumor volumes were monitored throughout the study in hu-PBMC-HNSCC mice injected peritumorally with EGFP-LNPs (n = 5) and control mice without LNP injection (no LNPs, n = 4). The experimental protocol for these experiments is outlined in Fig. 7A, and Materials and Methods. Data are presented as mean ± SEM. Welch's t-test was used to evaluate statistical differences between groups. (B) At the study endpoint (day 41), tumors were excised and their volumes were measured as described in the Materials and Methods. Tumor volumes from EGFP-LNP-injected and no LNP mice are shown as mean ± SD. Statistical significance was assessed using an unpaired t-test.

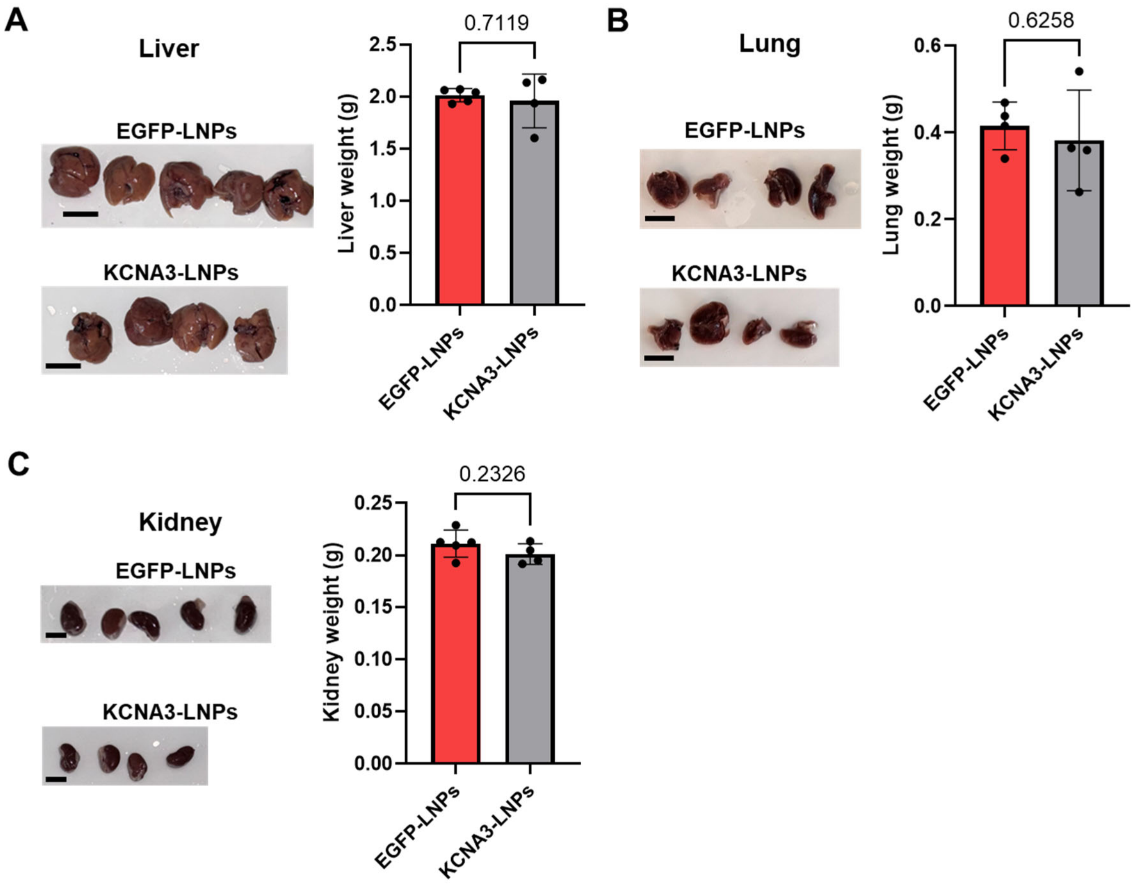


**Figure S10. Weights of the livers, lungs, and kidneys dissected from hu-PBMC-HNSCC mice injected with EGFP-LNPs or KCNA3-LNPs.** Images of the livers (A), lungs (B), and kidneys from hu-PBMC-HNSCC mice injected with EGFP-LNPs (n = 5) and KCNA3-LNPs (n = 4) are shown on the left, with a scale bar of 10 mm. The bar graph on the right displays the corresponding weights as the mean ± SD. The experimental protocol for these experiments is outlined in Fig. 7A, and the organs were dissected at the study endpoint. Statistical significance was assessed using an unpaired t-test.

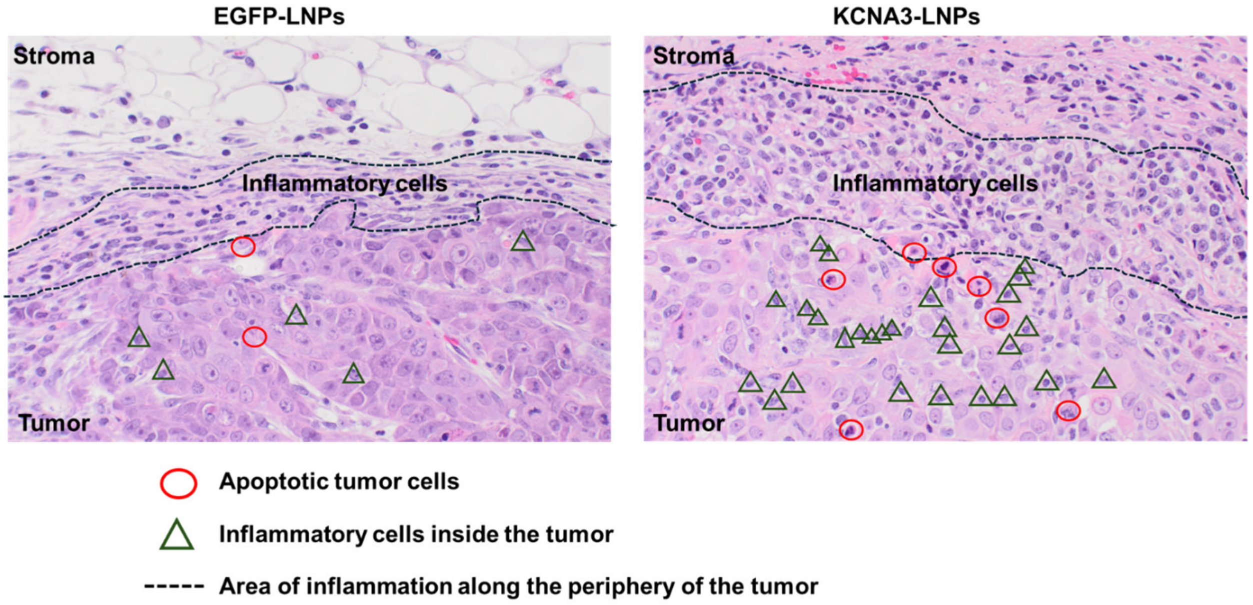


**Figure S11. Histopathological evidence of enhanced inflammatory infiltration and tumor cell death in hu-PBMC-HNSCC tumors treated with KCNA3-LNPs.** Representative hematoxylin and eosin (H&E)-stained sections of hu-PBMC-HNSCC tumors comparing treatment with EGFP-LNPs (left) to KCNA3-LNPs (right). Images were captured at a magnification of 40x. These are the same images shown in Fig. 8B, with detailed annotations by a board-certified pathologist indicating the tumor regions, inflammatory cell infiltrates, and apoptotic tumor cells. The area delimited by dashed lines highlights peritumoral regions characterized by a high density of inflammatory cells. Tumors treated with KCNA3-LNPs exhibited more prominent, thick bands of mononuclear inflammation at the tumor–stroma interface and within the immediate adjacent tumor parenchyma compared to the EGFP-LNP-treated controls. Red circles indicate apoptotic or degenerating tumor cells, identified by condensed or fragmented nuclei and cytoplasmic shrinkage, while green triangles denote representative inflammatory cells infiltrating the tumor periphery. These images are representative of multiple fields examined per group.

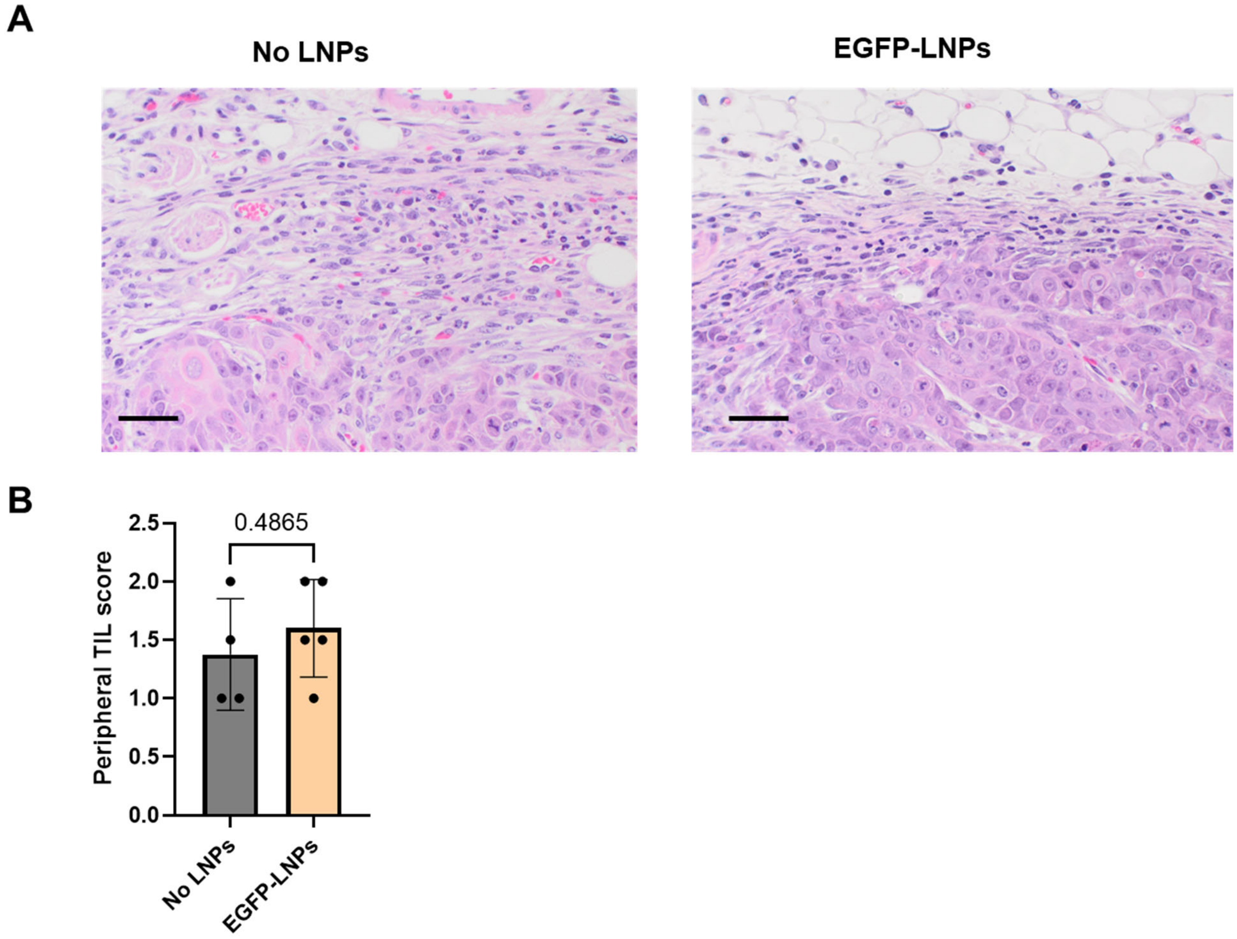


**Figure S12. Comparison of immune cell infiltration in EGFP-LNP-injected hu-PBMC-HNSCC mice and mice that did not receive LNPs.** (A) Representative H&E-stained tumor tissue from hu-PBMC-HNSCC mice not injected with LNPs (No LNPs) and mice treated with EGFP-LNPs, showing no significant differences in peripheral tumor inflammatory responses. Images at 40x magnification, Scale bar = 50 µm. (B) Peripheral TIL score was assessed by a board-certified pathologist (see Materials and Methods, and

Table S1) in KCNA3-LNPs (n= 4 mice) and EGFP-LNPs-treated (n= 5 mice) mice. The bars represent the mean ± SD, and significance was determined by Mann-Whitney U Test.

**Table S1 Semiquantitative histopathology analysis of intratumoral lymphocyte infiltration in hu-PBMC-HNSCC mice** Formalin-fixed, paraffin-embedded tumor sections from treatment groups in hu-PBMC-HNSCC humanized mice were stained with hematoxylin and eosin (H&E) using standard protocols. Blinded evaluation was performed by a board-certified pathologist. Immune cell infiltration was scored on a scale of 0 to 3, with 0 indicating minimal or no infiltration, 1+ indicating mild, 2+ indicating intermediate, and 3+ indicating severe infiltration. These scores represent the extent of immune cell presence across the entire tumor sections.

| **Group** | **Peripheral TILs** | **Central TILs** | **Peripheral Neutrophils** | **Central Neutrophils** | **Basophils** |
|---|---|---|---|---|---|
| KCNA3 #1 | 2-3+ | 1-2+ | 1+ | 1+ | 1+ |
| KCNA3 #2 | 2-3+ | 1-2+ | 1+ | 0-1+ | 1+ |
| KCNA3 #3 | 2+ | 1+ | 1+ | 1+ | 1+ |
| KCNA3 #4 | 2-3+ | 1+ | 1+ | 1+ | 1+ |
| EGFP #1 | 2+ (focal 3+) | 1-2+ | 1-2+ | 1+ | 0-1+ |
| EGFP #2 | 2+ | 1+ | 0-1+ | 1+ | 1+ |
| EGFP #3 | 1-2+ | 1+ | 1+ | 1+ | 2+ |
| EGFP #4 | 1+ (focal 2+) | 1+ | 1+ | 1+ | 2+ |
| EGFP #5 | 1-2+ | 1+ | 1+ | 1+ | 2+ |
| No NP #1 | 0-1+ | 0-1+ | 1+ | 1+ | 3+ |
| No NP #2 | 1-2+ | 1+ | 1-2+ | 1+ | 2+ |
| No NP #3 | 1+ | 1+ | 2+ | 1+ | 2+ |
| No NP #4 | 1+ (focal 2+) | 1+ | 2+ | 1+ | 2+ |
| No NP No PBMC #1 | 1+ | 0 | 2+ | 0 | 3+ |
| No NP No PBMC #2 | 1+ | 0 | 2+ | 0 | 3+ |

**Table S2 Antibodies used for LNP functionalization and Flow Cytometry experiments**

| Marker | Fluorophore | Clone | Vendor | Catalog # | RRID |
|---|---|---|---|---|---|
| Kv1.3 | N/A | N/A | Alomone | APC-101-GP | AB_2340958 |
| CD19 | APC-CY7 | HIB19 | Biolegend | 302218 | AB_314248 |
| CD3 | BV 421 | UCHT1 | Biolegend | 300434 | AB_10962690 |
| CD4 | BV 785 | OKT4 | Biolegend | 317441 | AB_2561365 |
| CD8 | BV 711 | SK1 | Biolegend | 344734 | AB_2565243 |
| CD56 | PE Dazzle 594 | HCD56 | Biolegend | 318347 | AB_2563563 |
| CD16 | PerCP | B73.1 | Biolegend | 360719 | AB_2563686 |
| CD14 | PE | 3G8 | Biolegend | 302007 | AB_314207 |
| CD69 | BV 711 | FN50 | Biolegend | 310943 | AB_2566465 |
| CD3 | BV 605 | OKT3 | Biolegend | 317321 | AB_11126166 |
| Zombie Red | N/A | N/A | Biolegend | 423109 | N/A |
| Zombie UV | N/A | N/A | Biolegend | 423107 | N/A |
| CD5 Biotin | N/A | UCHT2 | Biolegend | 300604 | AB_314090 |
| Biotin Mouse IgG1, κ Isotype Ctrl | N/A | MOPC21 | Biolegend | 400103 | AB_326427 |
